# DHHC7 palmitoylates KRAS4A and promotes mutant KRAS-driven pancreatic cancers

**DOI:** 10.64898/2026.03.31.715686

**Authors:** Wenzhe Chen, Seungmin Yeom, George Maio, Xiao Chen, Xuan Lu, Jiaqi Zhao, Neha Arora, Yinong Liu, Leah M. Ziolkowski, Martin Ian P Malgapo, Maurine E. Linder, Kay F. Macleod, Yong Zhou, Hening Lin

**Affiliations:** Department of Medicine and Department of Chemistry, The University of Chicago, Chicago, IL 60637, USA; Department of Chemistry and Chemical Biology, Cornell University, Ithaca, NY 14850, USA; Department of Diagnostic and Biomedical Sciences, School of Dentistry, University of Texas Health Science Center, Houston, TX 77054, USA; Ben May Department for Cancer Research, University of Chicago, Chicago, IL, USA, 60637; Department of Molecular Medicine, Cornell University, Ithaca, NY 14850, USA; Department of Integrative Biology and Pharmacology, McGovern Medical School, University of Texas Health Science Center at Houston, Houston, TX 77030, USA; Howard Hughes Medical Institute; Department of Medicine and Department of Chemistry; Center for Chemical Biology and Therapeutics; Ludwig Center at the University of Chicago; The University of Chicago Comprehensive Cancer Center; The University of Chicago, Chicago, IL 60637, USA

## Abstract

KRAS mutations underlie many human cancers. While inhibitors such as Sotorasib and Adagrasib targeting KRAS mutants have shown promise, additional strategies are required to address the broader spectrum of KRAS-driven cancers, particularly those displaying drug resistance. Thus, there is a need to better understand KRAS signaling and develop new therapeutic strategies. Here we show that KRAS4A is palmitoylated on Cys180 by a palmitoyl transferase, DHHC7 (gene name *ZDHHC7*). Palmitoylation promotes KRAS4A plasma membrane localization and importantly nanoclustering. This in turn promotes the activation of ARAF and RAF1, but not BRAF. DHHC7 and KRAS4A Cys180 palmitoylation are important for the normal and anchorage independent growth of pancreatic cancer cell lines. Depletion of *ZDHHC7* dramatically inhibits pancreatic tumor growth in mouse xenograft models. These studies provide new understandings about how palmitoylation regulates KRAS4A activity and suggest DHHC7 as a promising new target for KRAS mutant cancers.

## Introduction

Ras proteins, encoded by the HRAS, NRAS, and KRAS genes, are essential regulators of cell growth and proliferation, with mutations in KRAS contributing to nearly 20% of all human cancers and over 90% of pancreatic cancers.^1,2^ KRAS has long been considered ‘undruggable’ due to its lack of accessible binding pockets and its high affinity for GTP, making it difficult to inhibit effectively with traditional small molecules.^3^ Although recent breakthroughs in developing mutant KRAS inhibitors such as Sotorasib and Adagrasib have shown promising success, they face significant limitations, including the emergence of resistance and limited efficacy.^4,5^ These challenges underscore the need for innovative approaches, which requires a better understanding of how KRAS functions.

KRAS is alternatively spliced into two isoforms, KRAS4A and KRAS4B, with KRAS4B traditionally viewed as the major oncogenic driver. However, recent studies suggest a more significant role for KRAS4A than previously recognized.^6^ One study found that KRAS4A accounts for 10-50% of total KRAS expression, challenging earlier assumptions about its limited relevance. Another study showed that KRAS4A is more effective than KRAS4B in driving cellular transformation through the mitogen-activated protein kinase (MAPK) pathway, suggesting that KRAS4A plays a prominent role in tumorigenesis.^7,8^

The two isoforms primarily differ in their C-terminal hypervariable regions (HVR), where critical post-translational modifications (PTMs) occur. One key modification is S-palmitoylation, which regulates protein localization and protein-protein interactions; notably, KRAS4A undergoes S-palmitoylation on C180 while KRAS4B does not.^7,9,10^ Given KRAS4A’s superior ability to drive MAPK-mediated tumorigenesis compared to KRAS4B, *S*-palmitoylation may be a pivotal modification regulating tumor growth. Thus, identifying the specific palmitoyl transferases involved in KRAS4A S-palmitoylation may reveal new therapeutic targets for KRAS-driven cancers.

Here, we demonstrate that KRAS4A S-palmitoylation is important for its downstream MAPK pathway activation by influencing its membrane localization and nanoclustering or oligomerization, which is critical for the activation of its effector protein Raf in an isoform-specific manner. Furthermore, we identify DHHC7 as a major palmitoyl transferase for KRAS4A and regulator of the MAPK pathway. Reducing KRAS4A *S*-palmitoylation levels by depleting *ZDHHC7* significantly decreases tumorigenesis in KRAS mutant-driven pancreatic cancers suggesting DHHC7 could be a target for treating these cancers.

## Results

### *S-*Palmitoylation of KRAS4A is important for its RAF isoform specific interaction and downstream signaling

Mutations in KRAS4A can lead to dysregulation of the MAPK pathway, resulting in uncontrolled cell proliferation and cancer. To investigate the effect of KRAS4A *S*-palmitoylation on MAPK signaling, we ectopically expressed KRAS4A wild-type (WT) or mutants (C180S, G12D, and G12D/C180S) in HEK293T cells. G12D is the most common oncogenic version of KRAS and is more active than WT KRAS4A, while the C180S mutation replaces the palmitoylated cysteine with serine, and G12D/C180S is the double mutant. We used Western blotting to measure phosphorylated ERK (pERK) levels. KRAS4A stimulates ERK by binding to and activating the RAF kinases ARAF, BRAF, and RAF1. Therefore, we also measured RAF phosphorylation levels. As we expected, the C180S *S*-palmitoylation mutant decreases cellular pERK levels (Figure 1a). To our surprise, only the phosphorylation of RAF1 and ARAF, but not BRAF, was affected by KRAS4A *S*-palmitoylation (Figure 1b).

**Figure 1.**
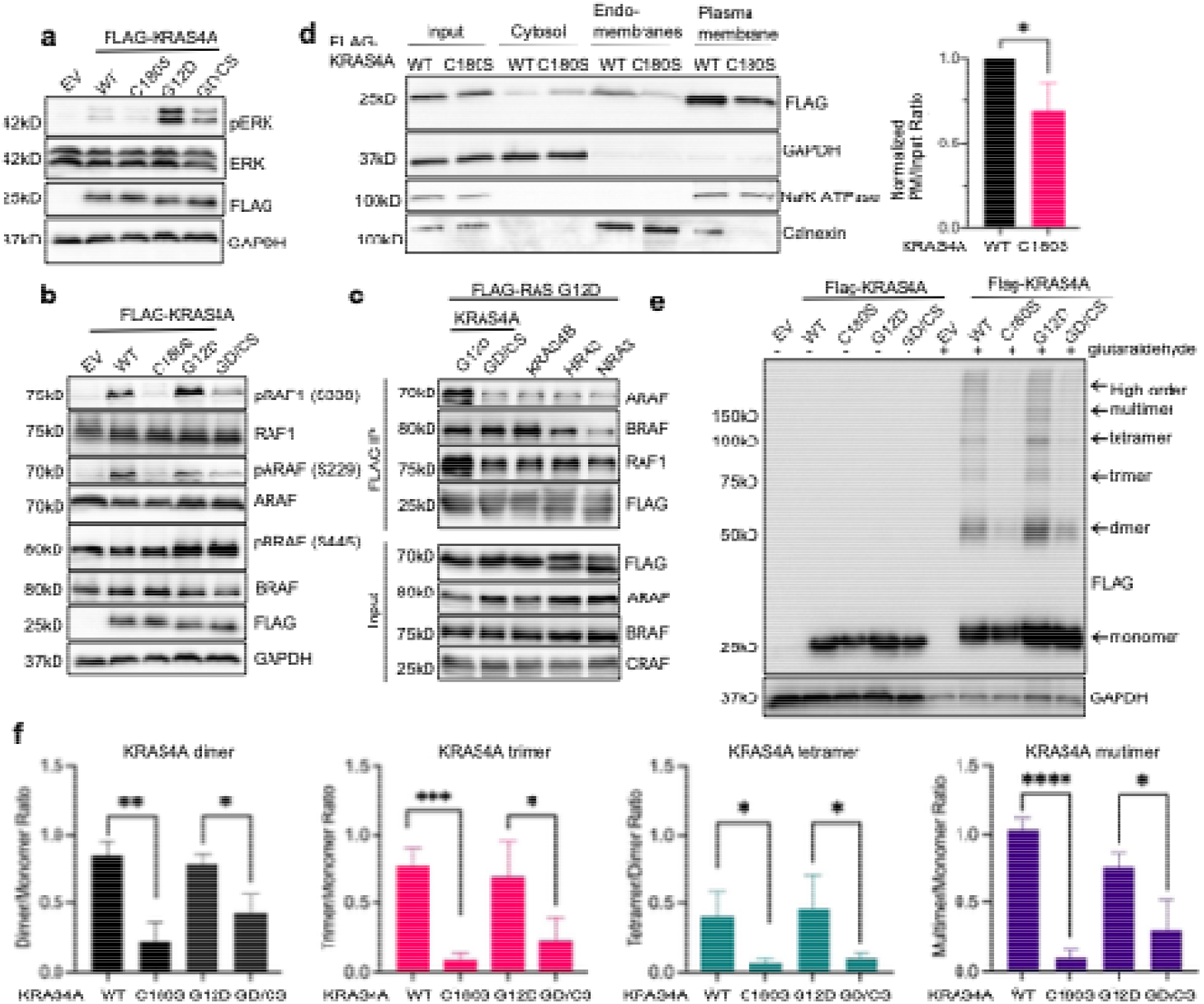
*S*-Palmitoylation is important for KRAS4A clustering, RAF isoform specific interaction, and downstream signaling. (**a**-**b**) Palmitoylation site mutation affects phosphorylation levels of ERK and different isoforms of RAF. HEK293T cells were transfected with FLAG-KRAS4A WT, C180S, G12D and G12D/C180S, pERK level (a) and phosphorylation levels of different RAF isoforms (b) were analyzed by western blot. (**c**) Palmitoylation affects KRAS4A interaction with different RAF isoforms. HEK293T cells were transfected with different RAS isoforms and their preferential interaction with different RAF isoforms was detected by FLAG IP. (**d**) Palmitoylation has a small effect on KRAS4A plasma membrane localization. HEK293T cells transfected with FLAG-KRAS4A WT and C180S were lysed and separated into cytosol, endo-membranes and plasma membrane fractions. The amount of FLAG-KRAS4A present in each fraction was detected and quantified using western blot. (**e-f**) Palmitoylation affects KRAS4A nanoclustering. Different KRAS4A constructs were expressed in HEK293T cells. The nanoclustering of KRAS4A was detected by intact cell glutaraldehyde crosslinking and western blot (e). Different oligomers were labeled based on the corresponding molecular weight and quantified in (f). The results in (d) and (f) are shown as mean ± SD of n = 3 independent western blot experiments. ns, not significant. \**P* < 0.05; \*\**P* < 0.01; \*\*\**P* < 0.001. Statistical significance was determined by two-tailed Student’s t-test for comparisons between two groups and by one-way ANOVA for comparisons among multiple groups.

The preferential activation of RAF isoforms by RAS proteins was previously reported for KRAS4B, HRAS and NRAS, but not KRAS4A.^11^ The author suggested that the preferential activation was regulated by the interactions between the RAS HVR and RAF N-termini. Consistent with this, replacing the HRAS HVR with that of KRAS4A or KRAS4B shifted RAF isoform binding accordingly, with the KRAS4B HVR chimera showing markedly reduced RAF1 and ARAF binding (Figure S1b). The positively charged polybasic regions on KRAS4B HVR increases its affinity towards the negatively charged BRAF N-terminus. Our data shown in Figure 1c and Figure S1a indicated that KRAS4A tends to have a stronger interaction with ARAF and RAF1 than with BRAF, and this specificity is dependent on C180 palmitoylation as the G12D/C180S mutant of KRAS4A behaves essentially like KRAS4B. Thus, C180 palmitoylation plays an important role in specifying the preferential activation of RAF1 and ARAF over BRAF by KRAS4A.

### *S-*Palmitoylation is important for KRAS4A nanoclustering

*S*-palmitoylation is well known to promote membrane localization^12,13^ and indeed it has been previously shown that S-palmitoylation of RAS affects its plasma membrane localization.^9^ Unlike HRAS and NRAS, which lose their plasma membrane localization completely after loss of S-palmitoylation, KRAS4A retains significant plasma membrane localization after C180S mutation. We used subcellular fractionation and confocal imaging to quantify the plasma membrane localization of KRAS4A WT and C180S mutant (Figures 1d, S1a and b). The C180S mutant retained >50% plasma membrane localization, which is also consistent with previous findings.^9^ Compared with the 80-90% decrease in pERK levels observed in Figure 1a and quantified in Figure S2c, the changes in KRAS4A plasma membrane localization may not fully explain the effect of S-palmitoylation. Consistent with this, in a fractionation experiment, pERK/ERK in the plasma membrane and endomembrane fractions of cells expressing C180S was only about 20% and below 10% of that in WT-expressing cells, respectively, even after normalization to the input level of each protein (Figure S3), indicating that the membrane-localized C180S mutant is not functionally normal.

Another important mechanism for KRAS4A activation is nanocluster formation. Although it has been a controversial topic over the past few years, the field has slowly reached a consensus that the HVR region of Ras protein is the driving force for the nanocluster formation.^14^ However, how HVR governs nanoclustering formation is not clear. We tested the effect of the C180S mutation on KRAS4A nanocluster formation using in-cell crosslinking followed by western blot. KRAS4A in a nanocluster will be crosslinked and appear as higher molecular weight species (dimer, trimer, tetramer, and other larger species) on the blot. Indeed, we observed these high molecular weight species. Furthermore, mutation of the *S*-palmitoylation site decreased nanoclustering of KRAS4A (Figure 1e, f), suggesting that KRAS4A C180 palmitoylation promotes its nanoclustering.

To further confirm the effect of the KRAS4A S-palmitoylation site mutation on KRAS4A nanoclustering, we detected KRAS4A molecules on plasma membrane sheets using electron microscopopy.^15^ The results suggest that the overall nanoclustering level decreases with the S-palmitoylation mutant for both KRAS4A WT and G12D (Figure S2d, e). Overall, our in-cell crosslinking western blot and plasma membrane sheet EM imaging support that S-palmitoylation promotes KRAS4A nanoclustering.

To test whether nanoclustering is driven by KRAS4A itself or induced by an effector, we transiently knocked down ARAF, which has been reported to form condensates at the plasma membrane ^16^, and repeated the crosslinking experiment. ARAF knockdown caused little change in the crosslinked KRAS4A species (Figure S4a), suggesting that S-palmitoylation does not promote KRAS4A nanoclustering through ARAF-mediated condensation. To ask whether the HVR is sufficient for clustering, we fused the KRAS4A HVR to monomeric mNeonGreen, which lacks the G-domain. The mNeonGreen-KRAS4A HVR alone formed crosslinked high-molecular-weight species, whereas the mNeonGree-KRAS4A C180S HVR formed far fewer (Figure S4b), indicating that the HVR is sufficient for clustering in a palmitoylation-dependent manner. The KRAS4B HVR also formed crosslinked species, so palmitoylation is not the only HVR feature that can support clustering in this assay.

### Chemically-induced clustering of S-palmitoylation deficient KRAS4A restores its Raf isoform specific interaction and downstream signaling

Given that S-palmitoylation affects both KRAS4A plasma membrane localization and nanoclustering, we wanted to determine whether nanoclustering is important for KRAS4A downstream signaling. Using the chemical induced dimerization system^17,18^ where we can artificially induce KRAS4A clustering by the addition of a small molecule, we first fused an FKBP domain to the N-terminus of KRAS4A G12D/C180S, the S-palmitoylation deficient mutant that has less nanocluster formation (Figure 2a). The FKBP domain will dimerize upon the addition of a small molecule, AP20187, possibly promoting KRAS4A nanoclustering. Since the induced dimerization happens in the absence of S-palmitoylation, it enables us to specifically examine the effect of nanoclustering on KRAS4A downstream signaling.

**Figure 2.**
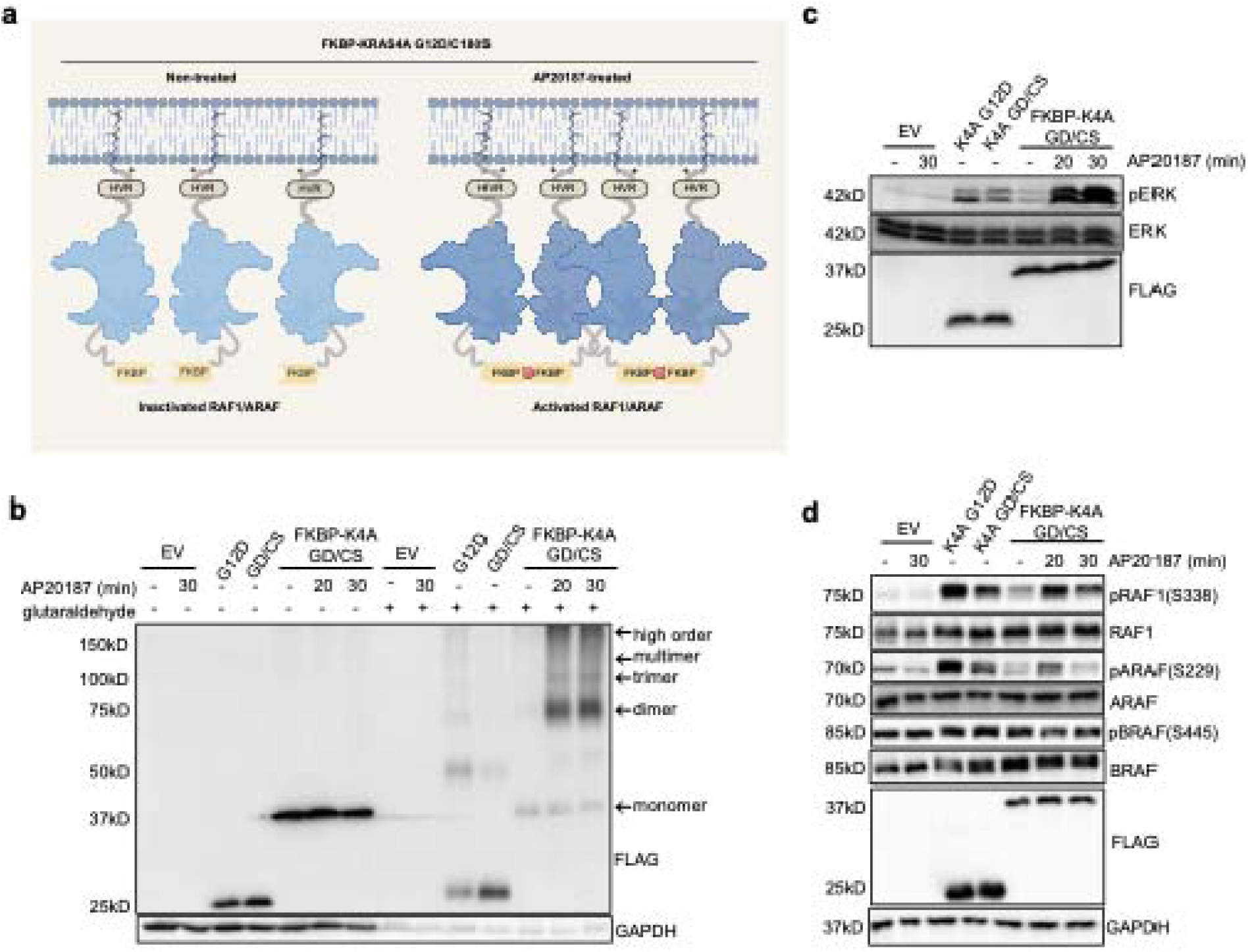
Chemical-induced clustering of *S*-palmitoylation deficient KRAS4A restores its Raf isoform specific interaction and downstream signaling. (a) Experiment design for chemical-induced FKBP-KRAS4A G12D/C180S nanoclustering. With the addition of the small molecule AP20187, FKBP-fused KRAS4A C180S would form nanocluster without S-palmitoylation. (b-c) Experimental validation of the chemical-induced nanoclustering of FKBP-KRAS4A G12D/C180S. HEK293T cells were transfected with different KRAS4A mutants and FKBP-fused KRAS4A G12D/C180S, and dimerization induced AP20187 was added for various amounts of time. After lysing the cells using 1% NP40, the nanoclustering of HEK293T KRAS4A was detected by intact cell glutaraldehyde crosslinking and western blot (b). Different oligomers were labeled based on the molecular weight. The pERK levels (c) and phosphorylation levels (d) of different RAF isoforms were analyzed by western blot.

We first show that this inducible dimerization system can induce not only dimers, but also higher order oligomers using in-cell crosslinking. In Figure 2b, upon AP20187 treatment, there was a significant increase of KRAS4A G12D/C180S dimer, trimer, and higher order oligomers compared to the non-treated group, supporting the inducible dimerization promotes nanoclustering. Thus, we successfully constructed an inducible nanoclustering system for the KRAS4A G12D/C180 mutant.

Then we measured the pERK level for FKBP-KRAS4A G12D/C180S before and after nanoclustering induction. AP20187 treatment strongly increased pERK signaling compared to untreated cells expressing FKBP-KRAS4A G12D/C180S (Figure 2c), suggesting that nanoclustering activates KRAS4A downstream signaling. AP20187 alone did not induce pERK in control HEK293T cells that do not express KRAS4A (Figure 2c), indicating that the ligand itself does not activate ERK signaling in these cells. AP20187-induced nanoclustering of FKBP-KRAS4A G12D/C180S was higher than natural nanoclustering of KRAS4A G12D (Figure 2b). This could explain why AP20187 induced higher pERK signals in cells expressing FKBP-KRAS4A G12D/C180S compared to KRAS4A G12D (Figure 2c).

Since we found that *S*-palmitoylation of KRAS4A differentially activates the three RAF isoforms, we investigated the effect of induced nanoclustering on RAF activation. Consistent with the finding that KRAS4A *S*-palmitoylation promotes ARAF/RAF1 activation but not BRAF activation, the induced nanoclustering only increased ARAF/RAF1 phosphorylation but not BRAF phosphorylation, suggesting that the nanoclustering of KRAS4A is critical for the activation of ARAF/RAF1 but not BRAF (Figure 2d). Overall, the inducible system indicated that nanoclustering promotes KRAS4A signaling and that beyond its effect on plasma membrane localization, S-palmitoylation also promotes KRAS4A signaling by promoting nanoclustering.

To further understand the interesting results between KRAS4A membrane localization and nanoclustering in a quantitative framework, we developed a simple thermodynamic model (Supplementary Note 1). The model links monomeric and dimeric KRAS4A, in the cytosol and on the membrane, through a closed thermodynamic cycle, such that at equilibrium any change that increases the membrane localization also increases the dimer fraction on the membrane. This model nicely explains how KRAS4A membrane localization and nanoclustering are linked and inter-dependent. In this framework, palmitoylation supplies a second lipid anchor and raises the membrane-bound fraction and thus increases dimerization (and oligomerization), whereas AP20187-induced dimerization of FKBP-KRAS4A G12D/C180S enters the same cycle by generating dimers directly. The model predicts that enforced dimerization increases membrane association of the FKBP-KRAS4A construct, as observed in fractionation experiments (Figure S5).

We next asked whether KRAS4A nanoclustering can be detected without overexpression. In-cell crosslinking of endogenous KRAS in MiaPaca-2 cells showed high-molecular-weight KRAS species (Figure S8). Because the available antibodies cannot reliably distinguish endogenous KRAS4A from KRAS4B in this assay, we used a pan-KRAS antibody, so this experiment reports total KRAS.

### DHHC7 mediated S-palmitoylation regulates KRAS4A clustering, Raf isoform specific interaction, and downstream signaling

S-palmitoylation is typically catalyzed by a group of enzymes containing conserved Asp-His-His-Cys catalytic residues, hence their names: DHHC proteins. ^19^ DHHC9 and its partner GCP16, work as the palmitoyl-transferase for HRAS and NRAS,^20^ but the palmitoyl-transferase for KRAS4A has not been reported. We performed a DHHC screen to identify the palmitoyl transferase responsible for KRAS4A palmitoylation. We overexpressed each of the 24 known mammalian DHHC proteins^21^ in HEK293T cells and measured KRAS4A S-palmitoylation levels using Alk14 metabolic labeling. After enriching for KRAS4A, we performed click-chemistry to add a fluorescent molecule to the alkyne group and detected S-palmitoylation by in-gel fluorescence. Our results suggested two potential palmitoyl-transferases for KRAS4A, namely DHHC3 and DHHC7 (Figure S6A). To rule out artifacts resulting from overexpression, we generated *ZDHHC3* and *ZDHHC7* CRISPR-knockout HEK293T cells. Using another assay to detect S-acylation, the Acyl-Biotin Exchange (ABE) assay, we found that either *ZDHHC3* or *ZDHHC7 KO* decreased KRAS4A acylation levels (Figure 3a). *ZDHHC7 KO* had a stronger effect than *ZDHHC3 KO* on KRAS4A acylation, suggesting that DHHC7 is the major palmitoyl-transferase for KRAS4A and thus DHHC7 is the focus of our studies moving forward.

**Figure 3.**
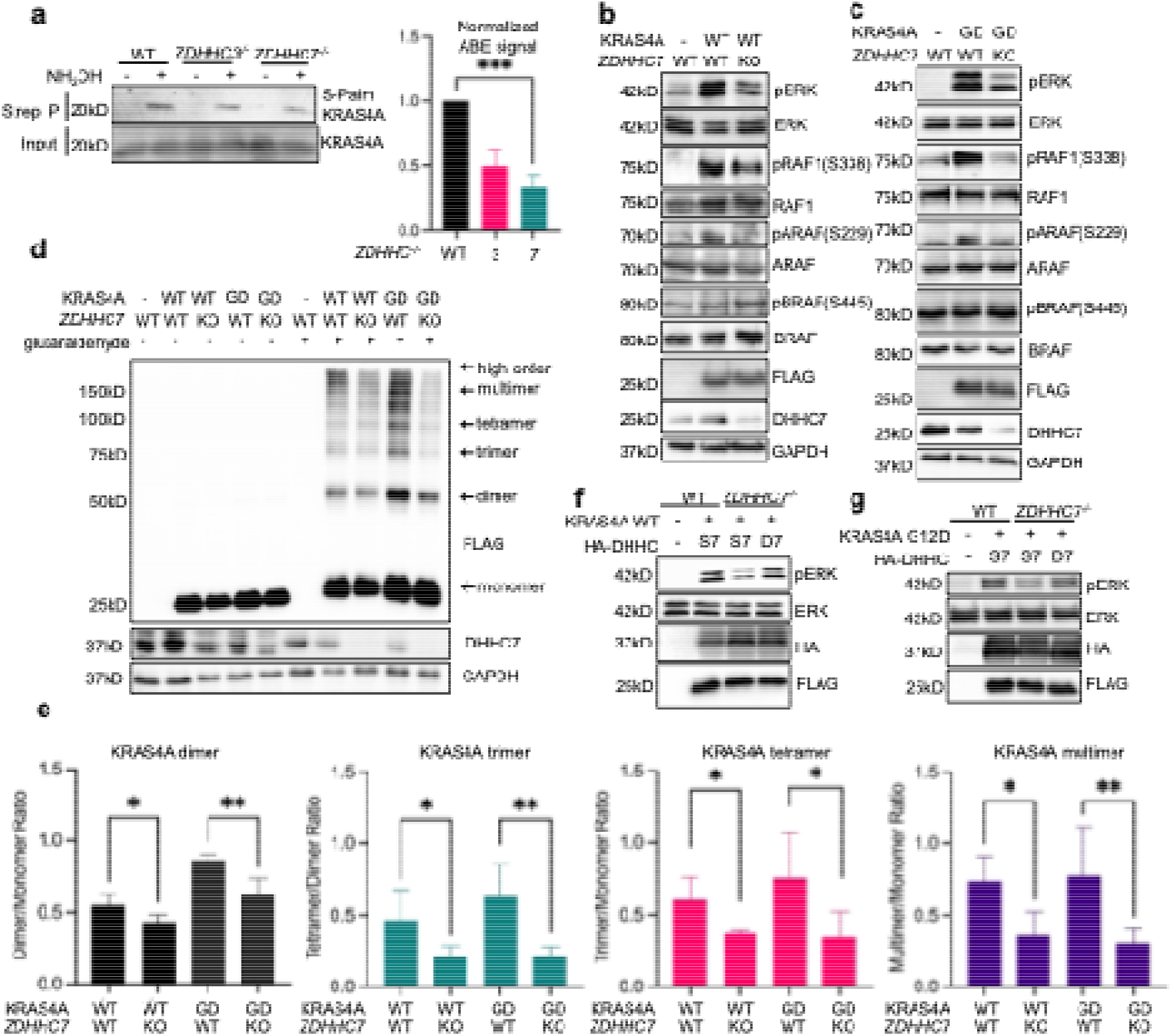
DHHC7 mediated *S*-palmitoylation regulates KRAS4A clustering, Raf isoform specific interaction, and downstream signaling. (**a**) DHHC7 and DHHC3 are KRAS4A palmitoyl transferases. *S*-Palmitoylation levels of endogenous KRAS4A were detected in HEK293T WT, *ZDHHC3* knockout, and *ZDHHC7* knockout cells using acyl-biotin exchange (ABE). Quantification was done using ImageJ. (**b, c**) DHHC7 affects the phosphorylation levels of ERK and RAF. FLAG-KRAS4A WT (**b**) or FLAG-KRAS4A G12D (**c**) was overexpressed in HEK293T WT and *ZDHHC7* KO cells. Phosphorylation levels of ERK and three different Raf isoforms were detected using western blot. (**d, e**) DHHC7 affects KRAS4A nanoclustering. In-cell crosslinking experiment was performed using KRAS4A WT and G12D overexpressed in HEK293T WT and *ZDHHC7* KO cells (**d**). The different oligomers of KRAS4A were labeled and quantified in (**e**). (**f**, **g**) The effect of DHHC7 on pERK is dependent on the catalytic activity of DHHC7. FLAG-KRAS4A WT (**f**) and G12D (**g**) were overexpressed in HEK293T WT and *ZDHHC7* KO cells with HA-DHHC7 or DHHS7 overexpression. The pERK levels were detected using western blot. The results in (**a**) and (**e**) are shown as mean ± SD of n = 3 independent western blot experiments. ns, not significant. \**P* < 0.05; \*\**P* < 0.01; \*\*\**P* < 0.001. Statistical significance was determined by two-tailed Student’s t-test for comparisons between two groups and by one-way ANOVA for comparisons among multiple groups.

To test the effects of *ZDHHC7* knockout on KRAS4A signaling, we transiently expressed KRAS4A WT or G12D mutant in *ZDHHC7* WT and KO HEK293T cells and blotted for pRAF and pERK. Compared to *ZDHHC7* WT cells, *ZDHHC7 KO* cells had decreased levels of pERK and pARAF/pRAF1, but not pBRAF (Figure 3b and 3c). To ensure that the effect we observed in *ZDHHC7 KO* cells was due to the lack of DHHC7 catalytic activity, we re-expressed DHHC7 or a catalytically inactive DHHS7 in the *ZDHHC7 KO* HEK293T cells. Overexpression of DHHC7 rescued pERK level, while the overexpression of DHHS7 did not (Figure 3f, 3g). These results are consistent with DHHC7 being a palmitoyl-transferase for KRAS4A and further confirms that S-palmitoylation promotes KRAS4A signaling.

Next, to determine whether *ZDHHC7 KO* affects KRAS4A nanoclustering, we expressed KRAS4A in *ZDHHC7* WT and KO HEK293T cells and used in-cell crosslinking to detect KRAS4A nanocluster formation. *ZDHHC7* KO cells exhibited less nanoclustering than WT cells (Figure 3d). We also quantified the change of different oligomers in Figure 3e and saw that different oligomers were decreased by the *ZDHHC7* KO in a pattern similar to that observed for the KRAS4A C180S mutant. Consistent with the partial effect of the C180S mutation on KRAS4A localization, KRAS4A remained at the plasma membrane in HeLa cells after ZDHHC7 knockdown, although more KRAS4A was observed on organelle membranes (Figure S7). These results support our hypothesis that DHHC7 regulates KRAS4A signaling by promoting its nanoclustering and plasma membrane localization.

In terms of downstream signaling, overexpression of DHHC7 increased pRAF1 and pERK in cells expressing WT KRAS4A but had little effect in cells expressing the palmitoylation-deficient C180S mutant (Figure S9a), indicating that the gain-of-function effect of DHHC7 on KRAS4A signaling depends on C180. Overexpression of DHHC3 and DHHC7 also increased KRAS4A S-palmitoylation (Figure S6) and the high-molecular-weight species in crosslinking assays, with a stronger effect from DHHC7 (Figure S9b). DHHC3 co-expression produced a smaller increase in pRAF1 than DHHC7 (Figure S9c), consistent with DHHC7 being the more effective enzyme for KRAS4A.

To test whether DHHC7 palmitoylates KRAS4A directly, we performed in vitro palmitoylation assays with purified DHHC7 and purified KRAS4A, KRAS4B, or HRAS. DHHC7 robustly palmitoylated KRAS4A, and most of the signal was removed by hydroxylamine, consistent with thioester-linked palmitoylation, whereas KRAS4B and HRAS showed little hydroxylamine-sensitive signal (Figure S10). This agrees with a previous in vitro study in which a peptide derived from the KRAS4A hypervariable region served as a substrate for purified ZDHHC7.^22^

### DHHC7 promotes anchorage-independent growth of KRAS4A G12D transformed 3T3 cells in an *S*-palmitoylation dependent manner

Given that palmitoylation of KRAS4A promotes its downstream signaling, we next investigated whether palmitoylation affects KRAS4A’s oncogenic properties. We generated 3T3 cells stably expressing pCDH empty vector, KRAS4A G12D, KRAS4A G12D/C180S, or KRAS4B G12D. Then we transiently overexpressed either DHHC7 or the catalytic dead mutant DHHS7 to compare the effect of KRAS4A *S*-palmitoylation on anchorage-independent cell growth in soft agar. Both KRAS4A G12D/C180S and KRAS4B G12D were used as *S*-palmitoylation deficient controls that we did not expect to be affected by DHHC7. The number of colonies formed was quantified using imageJ and CellProfiler (Figure 4a). Consistent with a previous report,^8^ KRAS4A transformed 3T3 cells had more colonies compared to 3T3 cells transformed with either KRAS4A G12D/C180S or KRAS4B G12D. This result suggests that KRAS4A *S*-palmitoylation is important for anchorage independent growth (Figure 4b). Compared to the overexpression of DHHS7, the overexpression of DHHC7 promoted the anchorage-independent growth of 3T3 cells transformed with KRAS4A G12D but not KRAS4A G12D/C180S or KRAS4B G12D (Figure 4c). Thus, DHHC7 promotes KRAS4A G12D’s transforming ability by promoting its *S*-palmitoylation. Consistent with these observations, DHHC7 overexpression also increased pERK signaling in 3T3 cells transformed with KRAS4A G12D, but not with KRAS4A G12D/C180S or KRAS4B G12D (Figure 4d).

**Figure 4.**
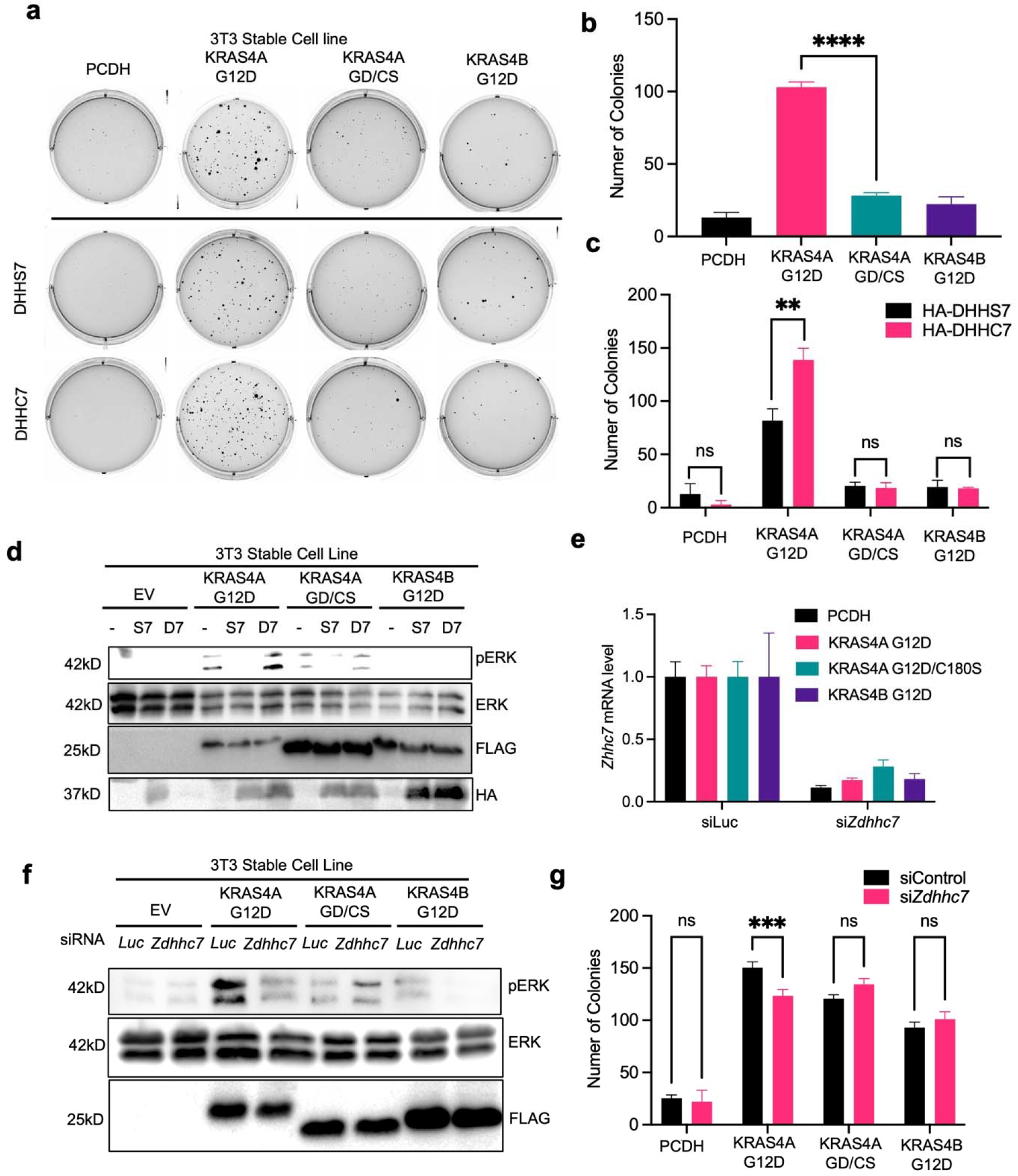
DHHC7 regulates anchorage-independent cell growth of KRAS4A-transformed NIH-3T3 cells in an *S*-palmitoylation-dependent manner. (**a-c**) NIH-3T3 were stably transformed with different KRAS mutants. Anchorage-independent cell growth of KRAS transformed NIH-3T3 cells were measured using soft-agar assay without or with the overexpression of HA-DHHS7 and HA-DHHC7 (**a**). The number of colonies formed was quantified in (**b**) and (**c**). (**d**) The pERK level of KRAS-transformed NIH-3T3 cells expressing no DHHC, HA-DHHS7, or HA-DHHC7 was analyzed by western blot. FLAG marks the KRAS4A and KRAS4B constructs, and HA marks DHHC7 and DHHS7. (**e-g**) Mouse *Zdhhc7* was knockdown in KRAS transformed 3T3 cells using siRNA. The mRNA level of *Zdhhc7* knockdown cells were measured using qPCR (**e**) the pERK level was detected using western blot (**f**), and the number of colony formation was measured and quantified (**g**). The results in (**b**), (**c**), (**e**) and (**g**) are shown as mean ± SD (n = 3). ns, not significant. \**P* < 0.05; \*\**P* < 0.01; \*\*\**P* < 0.001. Statistical significance was determined by two-tailed Student’s t-test for comparisons between two groups and by one-way ANOVA for comparisons among multiple groups.

We used siRNA to knock down mouse *Zdhhc7* in 3T3 cells transformed with different KRAS splice variants (Figure 4e). *Zdhhc7* knockdown decreased pERK level in KRAS4A G12D transformed 3T3 cells to a level similar to that of KRAS4A G12D/C180S or KRAS4B G12D transformed 3T3 cells (Figure 4f). This change in MAPK signaling was further validated in anchorage independent cell growth assay (Figure 4g). These results further confirmed the role of DHHC7 in promoting the transforming ability of KRAS4A.

### *ZDHHC7* knockdown inhibits mutant KRAS-driven pancreatic cancer in cellular and mouse models

KRAS is the driving oncogene in pancreatic ductal adenocarcinoma (PDAC) as 89% of human PDAC tumors carry activating KRAS mutations, with KRAS G12D being the most common. ^23^ Since we have shown that DHHC7-regulated S-palmitoylation is important for KRAS4A downstream signaling, we next investigated the effect of targeting DHHC7 for pancreatic cancer cell growth. We analyzed the differential expression level of *ZDHHC7* between normal pancreatic tissue and pancreatic tumor tissue using GEPIA web tools^24^ and found that *ZDHHC7* is expressed at higher levels in pancreatic tumor tissues compared to normal tissues (Figure S11a). We then analyzed the Kaplan-Meier survival curve for pancreatic cancer patients with low and high *ZDHHC7* expression levels using the Human Protein Altas database.^25^ The survival rate is higher for patients with low *ZDHHC7* expression compared to patients with high *ZDHHC7* expression (Figure S11b). This analysis is consistent with the hypothesis of *ZDHHC7* playing a role in PDAC progression. Interestingly, in pancreatic cancer patients, *ZDHHC7* and *KRAS* gene expression levels were highly correlated with a Pearson coefficient of 0.81 (Figure S11c).

To further examine the role of DHHC7 in pancreatic cancer, we used siRNA to knockdown *ZDHHC7* in various KRAS-driven cancer cell lines, including four KRAS mutant pancreatic cancer cell lines (MiaPaca-2, AsPC1, PANC0327 and Tu8988s) and one KRAS WT pancreatic cell line (BxPC3). We measured pERK levels and 2D cell proliferation 48-hr after *ZDHHC7* knockdown. *ZDHHC7* knockdown decreased pERK levels and inhibited 2D proliferation in KRAS mutant cell lines with the effect smaller in the KRAS WT cell line (Figure 5a and b). Given our data above shows that *ZDHHC7* KO also decreases pERK in cells expressing WT KRAS4A, the result that *ZDHHC7* knockdown also inhibits the KRAS WT pancreatic cancer cell line is not surprising. We also tested the effect of *ZDHHC7* knockdown in other types of cancer cell lines (lung and colon) harboring KRAS mutation (A549 and SW620) and found that *ZDHHC7* knockdown also decreased pERK and cancer cell growth (Figure S11d and S11e).

**Figure 5.**
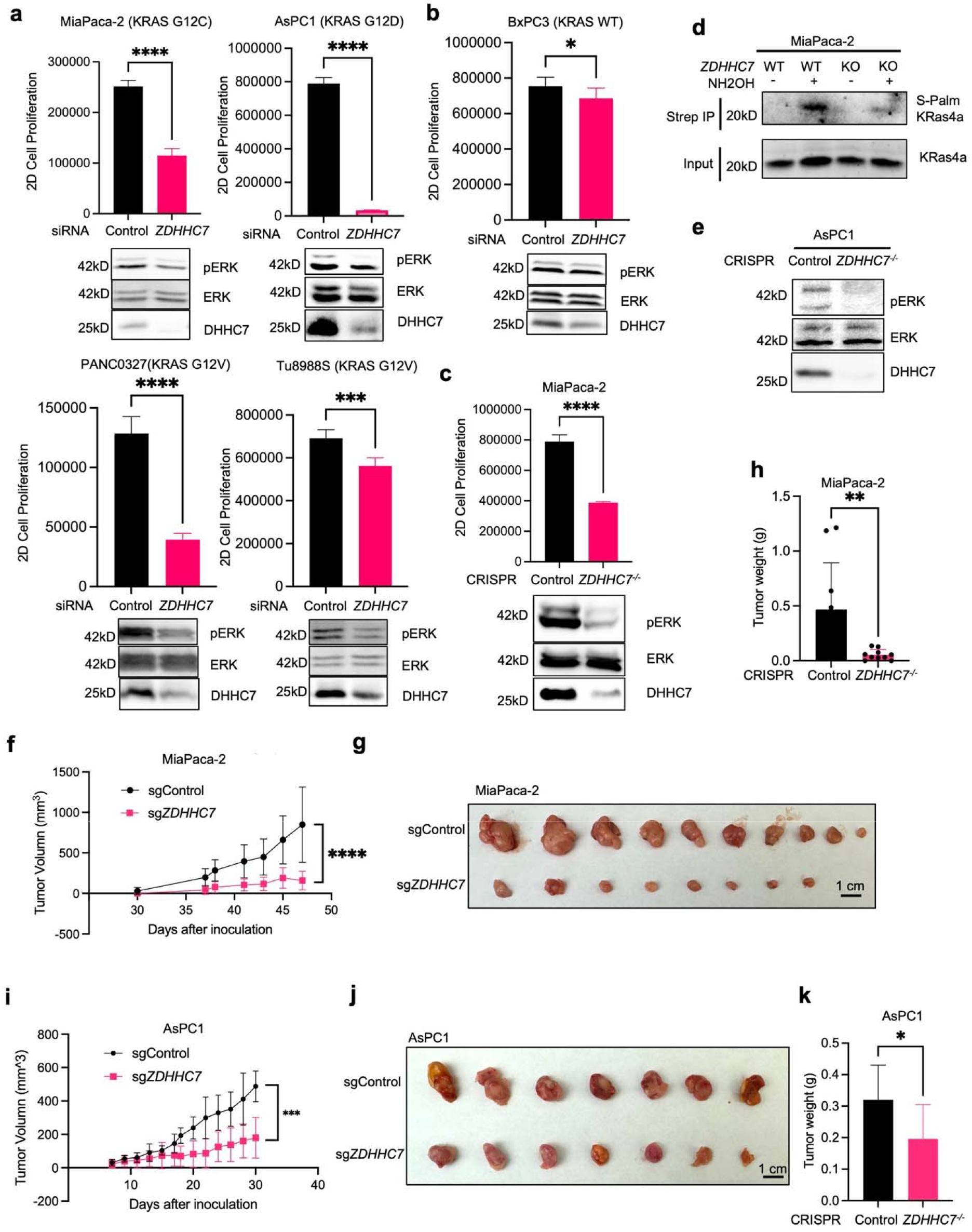
DHHC7 promotes KRAS-driven pancreatic tumor growth. (**a, b**) DHHC7 promotes pERK and 2-D cell growth of pancreatic cancer cells. *ZDHHC7* was knockdown using siRNA in four mutant KRAS-driven pancreatic cancer cell lines (AsPC1, MiaPaca-2, PANC0327 and Tu8988s) (**a**) and one non-KRAS driven pancreatic cancer cell line (BxPC3) (**b**). pERK was measured using western blot and 2-D cell growth was measured using Cell Titer Glow after 48-72 hrs. (**c-d**) *ZDHHC7* knockout affects pERK, 2-D cell growth, KRAS4A S-palmitoylation in MiaPaca-2 cell line. *ZDHHC7* was CRISPR knockout and pERK levels (**c**) were detected using western blot and 2D cell growth was measured after 72 hrs. The endogenous S-acylation level of KRAS4A was detected using ABE (**d**). (**e**) *ZDHHC7* was CRISPR knockout in AsPC1 cell line and pERK levels were detected using western blot. (**f-h**) *ZDHHC7* knockout suppresses MiaPaca-2 xenograft tumor growth. Pancreatic cancer xerograph was performed using MiaPaca-2 cells with control or *ZDHHC7* CRISPR knockout. The tumor volume was measured and quantified (f). The mice were dissected, and the final tumors were shown (**g**) and the tumor weights were quantified (**h**). (**i-k**) *ZDHHC7* knockout suppresses AsPC1 xenograft tumor growth. *ZDHHC7* CRISPR knockout was performed in AsPC1 cell line and similar xenograft experiment was performed. Tumor volume (i), tumors (j), tumor weight (k) are shown. Quantification data are shown as mean ± SD (western blot quantification, n = 3 independent experiments; cell proliferation, n = 8). ns, not significant. \**P* < 0.05; \*\**P* < 0.01; \*\*\**P* < 0.001. Statistical significance was determined by two-tailed Student’s t-test for comparisons between two groups and by one-way ANOVA for comparisons among multiple groups.

**Figure 6.**
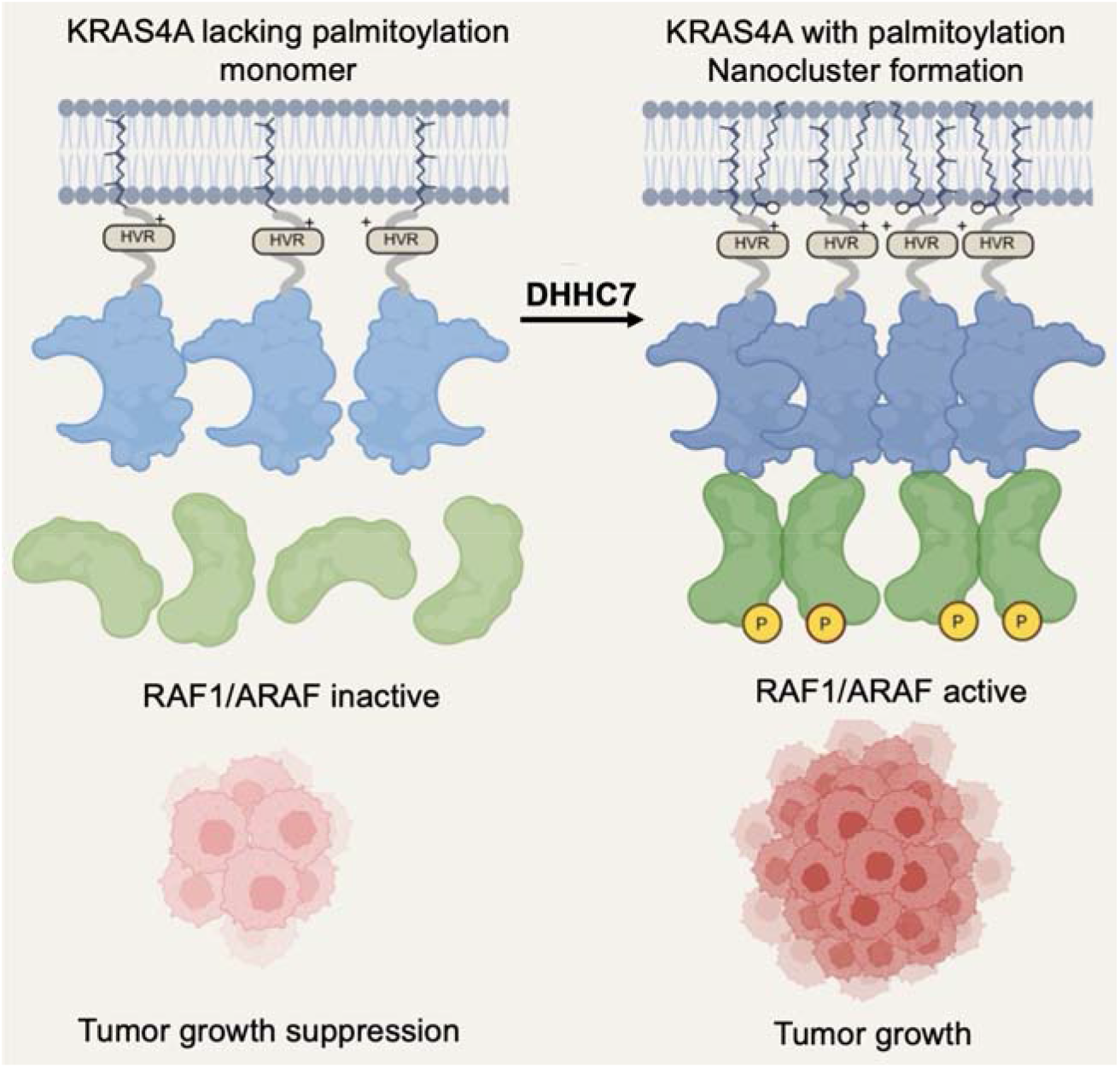
Proposed model of KRAS4A activation through DHHC7 mediated *S*-palmitoylation, which can activate RAF1/ARAF by formation of nanoclusters.

To examine the effect of DHHC7 on pancreatic cancer growth *in vivo*, we performed xenograft tumor studies using the MiaPaca-2 cell line in immune-compromised NSG mice. We first generated control and *ZDHHC7* CRISPR-KO MiaPaca-2 cell lines and verified knockout of *ZDHHC7*. The effect of ZDHHC7 KO on pERK level and 2-D cell growth was consistent with that of siRNA knockdown (Figure 5c and d). We performed ABE to detect endogenous KRAS4A S-palmitoylation levels by western blot. *ZDHHC7* knockout significantly decreased endogenous KRAS4A *S*-palmitoylation levels (Figure 5e). Next, we subcutaneously injected both the control and *ZDHHC7* CRISPR-KO MiaPaca-2 cell lines into the left (sgControl) and right (sg*ZDHHC7*) flank, respectively, of NSG mice to control for variations between animals. As expected, the sgControl MiaPaca-2 cells formed tumors by 45 days, but the *ZDHHC7* KO MiaPaca-2 cells failed to form tumors (Figure 5f-5h).

We further tested the effect of *ZDHHC7* knockout on xenograft tumor growth in vivo using two other pancreatic cancer cell lines, AsPC1 (Figure 5e and 5i–5k) and PANC0327 that carry the G12D and G12V mutations respectively (Figure S11f-h). Although we did not see as dramatic effects as in MiaPaca-2 xenograft, *ZDHHC7* knockout significantly decreased both AsPC1 and PANC0327 tumor growth *in vivo*.

ZDHHC7 is known to palmitoylate multiple substrates, including STAT3 and MAVS ^26,27^. To eliminate the effect of ZDHHC7 KO on its other substrates and test whether the effects of ZDHHC7 loss in pancreatic cancer can be mainly attributed to loss of KRAS4A palmitoylation, we re-expressed FKBP-KRAS4A G12D/C180S in ZDHHC7-KO MiaPaca-2 cells. ZDHHC7 KO reduced pERK and 2-D cell proliferation. Expression of the FKBP construct alone restored neither readout, whereas AP20187 treatment, which induces clustering independently of palmitoylation, restored pERK and restored proliferation to a level at least comparable to that of WT cells (Figure S12). Thus, in culture, the signaling and growth defects caused by ZDHHC7 loss are largely due to loss of KRAS4A palmitoylation-dependent clustering. We did not perform this rescue in xenografts due to technical challenges and therefore cannot exclude a contribution from other DHHC7 substrates to tumor growth in vivo. Our in vivo data using BxPC3 cell line, however, provide further evidence that the effect of ZDHHC7 knockout is mainly through regulating mutant KRAS. The BxPC3 pancreatic cancer cell line expresses WT KRAS and consistently, ZDHHC7 knockout in BxPC3 cells showed no significant difference in tumor volume compared to controls in the xenograft model (Figure S11i-k). The absence of any major tumor growth effect in a KRAS WT background provides independent in vivo evidence that the tumor-suppressing effect of ZDHHC7 knockout is mutant KRAS-dependent, rather than a general consequence of losing ZDHHC7 activity toward its other substrates. Thus, two independent lines of evidence converge on the same conclusion: restoring KRAS4A clustering alone is sufficient to rescue the signaling and proliferation defects of ZDHHC7 loss in culture (Figure S12) and removing mutant KRAS as a variable abolishes the effect of ZDHHC7 knockout on tumor growth in vivo (Figure S11i-k). Together these argue that mutant KRAS4A palmitoylation is the principal ZDHHC7-dependent driver in pancreatic cancer.

## Discussion

Our data here support a model that DHHC7 palmitoylates KRAS4A on Cys180 to promote its plasma membrane localization and nano-clustering, which in turn promotes downstream signaling through RAF1 and ARAF, but not BRAF. Depletion of DHHC7 inhibits the growth of human pancreatic cancer cell lines and dramatically inhibits pancreatic tumor growth in mouse xenograft models in vivo, suggesting DHHC7 is important for the growth of KRAS-driven pancreatic cancer cells and is a promising therapeutic target for pancreatic cancer.

*S*-palmitoylation has been well known to promote plasma membrane localization of peripheral membrane proteins like HRAS and NRAS ^10,28–31^. Our data suggest that for KRAS4A, the role of S-palmitoylation is not simply to deposit the protein at the plasma membrane. We showed that DHHC7 catalyzed *S*-palmitoylation on KRAS4A affects its ability to form nanoclusters. The importance of the nanocluster-promoting role for KRAS4A *S*-palmitoylation in downstream signaling was further demonstrated using a chemical-induced clustering system with the FKBP-KRAS4A G12D/C180S protein. We noted two things indicating that the amount of KRAS4A at the plasma membrane does not by itself account for the role of *S*-palmitoylation. First, a significant portion of KRAS4A is still localized to the plasma membrane without palmitoylation. Second, KRAS4A is reported to be more active towards ARAF when it is localized on intracellular membranes.^9^

In fractionation experiments, C180S remained associated with the plasma membrane and endomembrane fractions but showed much lower pERK, even after normalization to its expression level (Figure S3), indicating that the membrane-localized mutant is not functionally normal. At the same time, our thermodynamic model (Supplementary Note 1) indicates that membrane localization and oligomerization are not independent variables but two arms of a single closed cycle: because a KRAS4A dimer carries more membrane anchors than a monomer, localization makes oligomerization more favorable, and the oligomers so formed hold the membrane more tightly, so the two processes reinforce each other. Palmitoylation supplies the second lipid anchor and therefore acts as a physiological entry point into this loop, raising the membrane-bound fraction and thereby enriching dimers on the membrane, whereas a palmitoylation-deficient protein is retained on the membrane by its farnesyl anchor alone and enters the loop only weakly; enforced dimerization enters the same cycle from the other arm, generating dimers directly, and accordingly increases the membrane association of the FLAG-FKBP-KRAS4A G12D/C180S construct in fractionation experiments (Figure S5). Thus, membrane localization and nanoclustering are coupled processes that cannot be separated in cells, and we do not claim that nanoclustering acts independently of localization.

Several caveats apply to the chemical-induced clustering system. AP20187 alone did not induce pERK in cells lacking exogenous KRAS4A (Figure S9), but we cannot exclude that the FKBP fusion or AP20187-induced dimerization may affect the normal function of KRAS4A. This may be the reason why forced dimerization only partially recapitulates native palmitoylation-driven activation of RAF1 and ARAF (Figure 2d). The extent of rescue by forced dimerization also differs between readouts and cell systems: pERK and proliferation in ZDHHC7-KO MiaPaca-2 cells were restored to WT levels or higher (Figure S12). We also note that the transient phosphorylation of ARAF and RAF1 while pERK remains elevated (Figure 2d) is consistent with ERK-dependent negative feedback on RAF.^32–34^

Many integral membrane proteins are known to be regulated by S-palmitoylation and for these proteins, it is often proposed that *S*-palmitoylation promotes their lipid raft association.^13^ A recent report on MAVS palmitoylation raised a different model: *S*-palmitoylation helps the activated and aggregated MAVS to remain on the mitochondrial outer membrane.^27^ KRAS4A S-Palmitoylation as reported here may represent yet another example with S-palmitoylation promoting the nanocluster/aggregation of KRAS4A, which is important for its function. This makes sense especially because the downstream proteins, like ARAF, are also known to form phase condensates.^16^ It is easy to imagine that the aggregated KRAS4A may bind the aggregated ARAF much better than the non-aggregated counterparts due to a multivalence effect. It is not clear whether the KRAS4A nanocluster localizes to lipid rafts or not; even if it does localize to lipid rafts, it would only engage the inner leaflet of the membrane bilayer, making it different from the S-palmitoylation of integral membrane proteins. Another recent study suggested that without S-palmitoylation, KRAS4A localizes to mitochondria and interacts with HK1^35^. Interestingly, it was recently reported that KRAS4A preferentially associates with dual-saturated and dual-unsaturated phosphatidylserine (PS) species. This is distinct from KRAS4B, which heavily favors mixed-chain lipid species.^36^ This suggests that KRAS4A may prefer to line the boundaries of lipid rafts, where KRAS4A interacts with both the saturated lipids in the raft and the non-raft unsaturated lipids. This is similar to the domain boundary-favoring NRAS.^37^ This illustrates that while in general *S*-palmitoylation promotes membrane targeting, the specific effects and the mechanistic details of its function can be diverse.

Another novel mechanistic insight reported here is the differential activation of Raf isoforms by KRAS4A nanocluster. First, Raf proteins were known to be an important effector protein of Ras, but the differential functions of the three main Raf isoforms is not well studied. Here we show that KRAS4A preferentially activates ARAF and RAF1 but not BRAF (Figure 1e). Interestingly, KRAS4B was reported to preferentially activate BRAF due to the interaction between KRAS4B’s positively charged C-terminal and BRAF’s negatively charged N-terminus.^38^ It has also been suggested that RAF1 and BRAF differ in their ability to associate with PS lipids ^39^. Our findings here suggest that while positive charges at HVR promote BRAF activation, the S-palmitoylation-mediated nanoclustering promotes ARAF and RAF1 activation. This is further supported by the observation that the palmitoylation deficient KRAS4A C180S mutant activates BRAF better than ARAF and RAF1. The mechanism underlying this selectivity is not fully resolved. Forced dimerization of the palmitoylation-deficient mutant restores signaling through ARAF and RAF1 but not BRAF in the absence of the lipid modification, which favors a model in which the clustered state of KRAS4A, rather than a direct palmitoyl-dependent contact between the HVR and ARAF, determines which RAF isoforms are engaged. A recent ITC study found comparable G-domain affinities of KRAS4A and KRAS4B for RAF1 and proposed that isoform differences instead increase the population of KRAS-RAF1 complexes on the membrane [6], which is also consistent with an effect of membrane localization and nanoclustering.

Nanoclustering of KRAS has been shown to be important for its downstream signaling activation, but the mechanistic details of nanoclustering are still lacking ^40^. One key controversial point concerns the identity of the driving force for nanocluster formation. One proposal is that the dimeric interactions between two Ras molecules promote nanoclustering.^41^ Another point of view is that the interaction between the C-terminal HVR and lipids in the membrane is important.^42^ By showing that *S*-palmitoylation on HVR is critical for nanoclustering, our data support the importance not only of HVR, but specifically S-palmitoylation, in driving nanoclustering (Figure 1b). How *S*-palmitoylation affects KRAS4A nanoclustering is an interesting question. Our analysis in Supplementary Note 1 suggests that KRAS4A’s membrane localization and oligomerization (and by extension, nanoclustering) are thermodynamically linked, and thus, increasing KRAS4A membrane localization will increase its nanoclustering. This analysis provides a logical explanation for how S-palmitoylation increases KRAS4A nanoclustering: rather than acting on clustering directly, the palmitoyl group raises the membrane affinity of KRAS4A and so feeds the self-reinforcing loop between localization and oligomerization. This localization-driven self-assembly is not unique to KRAS4A and should apply to many other membrane-targeting proteins, derived from equilibrium thermodynamics (Supplementary Note 1).

The KRAS4A nanoclustering could further help to transduce signals from upstream to downstream partners. This is because upstream and downstream partners (e.g. receptor tyrosine kinases and ARAF) are also reported to form condensates^43–45^ ^16^. Our data supports that KRAS4A can form nanocluster by itself independent of ARAF condensates as we shown ARAF knockdown caused little change in KRAS4A crosslinking (Figure S4a) and mNeonGreen fused to the HVR of KRAS4A formed palmitoylation-dependent crosslinked species without the G-domain that binds to ARAF (Figure S4b). However, it is likely that the KRAS4A nanocluster can bind to the ARAF condensates more strongly due to multivalence effects, thus promoting downstream signaling. Nanoclustering and phase condensation may be related manifestations of locally concentrated KRAS proteins detected by different techniques, although they are not necessarily equivalent, and distinguishing between them, as well as characterizing the reversibility and dynamics of the clusters, is beyond the scope of this study.

Mutant KRAS underlies ∼20% of all human cancers. There has been a longstanding interest in targeting KRAS to treat human cancers and several inhibitors targeting mutant KRAS are approved.^46^ Our study here suggests a new way to treat KRAS mutant-driven cancers by targeting its palmitoyltransferases, DHHC7 and likely also DHHC3. Initially we were unsure about how big an effect would be observed in mutant KRAS pancreatic cancer as DHHC7 in principle should only affect KRAS4A, but not KRAS4B. However, the mouse xenograft experiments showed a strong effect of *ZDHHC7* depletion on tumors formed from PDAC cell lines expressing mutant KRAS. There are two possible reasons for this. One is that mutant KRAS4A is the major tumor driver as it can activate ARAF and RAF1 much better than KRAS4B ^8^. KRAS4A is reported to be important for cancer stem cell generation.^47^ It is possible that by affecting KRAS4A palmitoylation, *ZDHHC7* knockout pancreatic cancer cells cannot sustain normal tumor development. Given the distinct interactomes and regulation mechanisms of KRAS4A and KRAS4B previously reported ^8,48^, it makes sense that each splicing isoform governs different critical pathways in tumorigenesis. Another possibility is that DHHC7 also works on other substrate proteins that promote tumor growth, such as STAT3.^26^ It has been suggested that KRAS and STAT3 work together to promote tumorigenesis and STAT3 becomes the main tumor driver after KRAS inhibition.^49,50^ Thus, DHHC7 inhibition could provide a unique approach to target KRAS and STAT3 simultaneously and produce strong anticancer effects in pancreatic cancer. However, our cell line study (Figure S12) indicates that, in cell culture, the signaling and growth defects caused by *ZDHHC7* loss are largely due to loss of KRAS4A palmitoylation. Furthermore, in KRAS WT pancreatic tumor xenograft, the effect of *ZDHHC7* knockout is very minor, arguing that even in vivo, the effect of *ZDHHC7* knockout on tumor growth is mainly through regulating mutant KRAS4A.

Another observation of note is that the effect DHHC7 deficiency is much more pronounced in vivo in the xenograft mouse models than in 2-D or anchorage-independent growth in culture. This may be explained by the hypoxic nature of PDAC tumors in vivo compared to growth in culture at 20% oxygen;^51^ and we note that hypoxia promotes *ZDHHC7* expression and activity ^52^ consistent with DHHC7-mediated KRAS4A palmitoylation being more pronounced in vivo due to the hypoxic conditions. In summary, our work here suggests that DHHC7 is a promising candidate target for treatment of KRAS-driven pancreatic cancer, although our evidence is based on genetic depletion of DHHC7 and not on pharmacological inhibition. Developing DHHC7 inhibitors, therefore, will be an important direction to pursue in the future.

## Materials and Methods

### Mice

NOD scid gamma (NSG) mice NOD.Cg-Prkdc^scid^ Il2rg^tm1Wjl^/SzJ (RRID:IMSR_JAX:005557) were purchased from The Jackson Lab and then bred in house. All the mice were housed in a pathogen-free facility. The mouse protocols were approved by Cornell University’s Institutional Animal Care and Use Committee and University of Chicago’s Institutional Animal Care and Use Committee.

### Antibodies, reagents, and plasmids

Antibodies were purchased from commercial source including antibodies from commercial sources include phospho-p44/42 MAPK (ERK1/2) (Thr202/Tyr204) (CST, 4370, 1:1000 dilution), p44/42 MAPK (ERK1/2) (CST 9102, 1:100 dilution for immunofluorescence), Flag-HRP (Millipore A8592, 1:5000 dilution), phospho-RAF1 (Ser338) (CST 9427, 1:1000 dilution), phospho-BRAF (Ser338) (CST 2696, 1:1000 dilution), phospho-ARAF (Ser338) (CST 4431, 1:1000 dilution), β-Actin (C4) (Santa Cruz sc-47778, 1:1000 dilution), GAPDH-HRP (Santa Cruz sc-47724, 1:1000 dilution), HA-Tag (CST 3724, 1:1000 dilution), anti-mouse IgG HRP (CST 7076S, 1:2500 dilution), anti-rabbit IgG HRP (CST 7074S, 1:2500 dilution), DHHC7 (Abcam A138210, 1:1000 dilution). KRAS4A specific antibody was kindly provided by Dr. Mark R. Philips.

Flag-KRAS4A and FLAG-KRAS4B plasmids were obtained from Origene and mutated versions were generated using site-direct mutagenesis. The gene encoding FLAG-KRAS4A G12D, G12D/C180S and FLAG-KRAS4B G12D was cloned into PCDH empty vector with a CMV promoter for lentivirus preparation. The plasmid encoding Cas9 and DHHC7 sgRNA with CMV promoter for lentivirus preparation was obtained from Genecopoeia (Cas9: CP-LvC9NU-01, sgRNA: HCP257256-LvSG03-3-B). Murine ZDHHC1-23 plasmids were kindly provided by Dr. Masaki Fukata. Human and mouse DHHC7 siRNA was purchased from Santa Cruz (mouse: sc-155507, human: sc-93249).

### Chemical and biochemical regents

Glutaraldehyde (Millipore Sigma, 70% in H_2_O), Tetramethylrhodamine-azide (TAMRA-azide, 7130, Lumiprobe), tris[(1-benzyl-1H-1,2,3-triazol-4-yl)methyl]amine (TBTA, T2993, TCI Chemicals), CuSO_4_ (TCI Chemicals), tris(2-carboxyethyl)phosphine hydrochloride (TCEP, Sigma-Aldrich C4706), 2-bromopalmitic acid (2-BP, Sigma-Aldrich 21604), polyethylenimine hydrochloride (PEI, Polysciences), protease inhibitor cocktail (Sigma-Aldrich P8340), phosphatase inhibitor cocktail (Sigma P0044), universal nuclease (Thermo Fisher 88700,), dithiothreitol (Goldbio DTT100), Pierce ECL Western blotting substrate (Thermo Fisher), Clarity Max Western ECL substrate (Bio-Rad, 1705062), high-capacity streptavidin agarose (20359, Thermo Fisher), anti-Flag agarose gel (Sigma-Aldrich A2220,), hydroxylamine (50% solution in water, Sigma-Aldrich), N-ethylmaleimide (NEM, Sigma-Aldrich), Nitroblue tetrazolium chloride (Sigma-Aldrich) and CellTiter-Glo® 2.0 Cell Viability Assay (Promega). 15-Hexadecynoic acid (Alk14) was synthesized in-house.

### Cell culture, transfection, and co-immunoprecipitation

HEK293T, NIH 3T3, MiaPaca-2, BxPC3, PANC0327 and Tu8988S were cultured in Dulbecco’s modified Eagle’s medium (DMEM, Gibco) containing 10% fetal bovine serum (Gibco) and 1% antibiotic-antimycotic (Gibco). All the cell lines were purchased from ATCC. DHHC7^-/-^HEK293T and MiaPaca-2 cells were generated using CRISPR/Cas9 and verified using western blot and qPCR. NIH 3T3 cells stably expressing PCDH empty vector, KRAS4A G12D, KRAS4A G12D/C180S and KRAS4B G12D were generated using lentivirus transduction.

For the transient transfection, HEK293T cells were seeded one day prior to transfection in 10-cm dishes containing DMEM media with 10% FBS. When cells reach ∼70% confluency, plasmids encoding the indicated proteins were diluted in serum free DMEM, and polyethylenimine hydrochloride (PEI, Polysciences) was added at a DNA to PEI ratio of 1:3. The DNA-PEI mixture was incubated at room temperature for 30 minutes and then added dropwise to the HEK293T dishes. After 24 hours, cells were collected or treated for further analysis.

For siRNA transfection in NIH 3T3 and pancreatic cancer cell lines, Lipofectamine™ RNAiMAX (Invitrogen) was used according to the manufacturer’s instructions. Briefly, cells were seeded in 6-well plates the day before. When the cell reaches ∼70% confluency, 30 pmol RNAi duplex was diluted into 150 μL of Opti-MEM I reduced serum medium, and 9 μL of transfection agent was diluted into another 150 μL of Opti-MEM I reduced serum medium. The diluted RNAi duplex and transfection agent were combined and incubated at room temperature for 15 minutes before being added drop-wise to the wells. Cells were incubated for 36–48 hours and then treated for further analysis.

For co-immunoprecipitation (Co-IP) of Flag-KRAS4A with endogenous Raf isoforms in HEK293T cells, transfected cells were lysed with 1% NP-40 lysis buffer (25 mM Tris-HCl pH 8.0, 10% glycerol, 150 mM NaCl, 1% NP-40) containing protease inhibitor cocktail (Sigma, P8849) on ice for 30 mins with frequent vortexing. Flag-KRAS4A was then immunoprecipitated using anti-FLAG M2 agarose beads (Sigma). The lysate and IP samples were then analyzed by SDS-PAGE and western blot.

### Lentivirus transduction

For 3T3 cell stable cell line generation, lentivirus was generated by transfecting HEK293T cells with pCDH vector containing KRAS4A G12D, KRAS4A G12D/C180S and KRAS4B G12D along with packaging vector psPAX2 and envelope vector pMD2.G. After 48 hours, the cell culture media containing lentiviral particles was collected and filtered through 0.45 μm filters. The filtered virus was then used to infect 3T3 cells using 8 μg/mL polybrene. The cells were incubated with virus for 24 hrs before replacing media containing 5 mg/mL of puromycin. The cells were cultured with selection marker for a few passages and verified using western blot.

To knock out DHHC7 in MiaPaca-2 cells, we first generated MiaPaca-2 cells stably expressing Cas9 nuclease. Similarly, we generated lentivirus by transfecting HEK293T cells with a vector encoding Cas9 nuclease (GeneCopoeia), along with the packaging vector psPAX2 and envelope vector pMD2.G. After transduction, the cells were selected using 400 ng/mL G-418 for 48 hours and surviving cells were expanded and verified for Cas9 expression. Subsequently, second round of lentivirus transduction delivered sgRNA into the cells. Lentivirus containing sgRNA were generated by transfecting vectors expressing sgRNA targeting DHHC7 (GeneCopoeia) in HEK293T cells along with psPAX2 and pMD2.G. The resulting filtered cell culture media containing lentiviral particles was used to transduce MiaPaca-2 Cas9 cells. Infected cells were selected using 5 μg/mL puromycin for 48 hours and DHHC7 knockout was verified by immunoblotting and Q-PCR.

### In-cell crosslinking

Pre-treated HEK293T cells were collected into a 1.5 mL tube using cold PBS and split into two tubes. One third of the cells were used as control group without crosslinker treatment and 2/3 of cells were used for crosslinker treatment. The treated cells were resuspended in 3 mM glutaraldehyde solution with PBS and incubated at room temperature for 15 mins followed by 10 mins quenching using 3 mM Tris pH 7.4. The resulting cell pellets were washed again using PBS before lysing with 1% NP-40 lysis buffer (25 mM Tris-HCl pH 8.0, 10% glycerol, 150 mM NaCl, 1% NP-40, 1X protease inhibitor). The protein lysate was analyzed using SDS-PAGE and corresponding oligomerization state of protein of interest was detected based on molecular weight on western blot.

### Chemical-induced dimerization system

The cells were transfected with FKBP-fused plasmid. After 24 hrs, the cells were treated with DMSO or AP20187 for specific time to induce dimerization and then collected for in-cell crosslinking and western blot.

### In-gel fluorescence for S-palmitoylation detection using Alk14

HEK293T cells were transfected with Flag-KRAS4A and ZDHHCs-HA using PEI for 24 hours and treated with 50 μM Alk14 for 6 hours before collection. After washing, cells were lysed using 1% NP-40 lysis buffer (25 mM Tris-HCl pH 8.0, 10% glycerol, 150 mM NaCl, 1% NP-40). Flag-KRAS4A was purified using anti-Flag agarose beads, and a click chemistry mixture was added to each sample. This mixture contained 40 μL IP wash buffer (25 mM Tris-HCl pH 8.0, 150 mM NaCl, 0.2% NP-40), 2 μL of 2 mM TAMRA-azide, 2.4 μL of 10 mM TBTA, 2 μL of 40 mM CuSO_4_, and 2 μL of 40 mM TCEP. The mixture was vortexed and incubated in the dark at room temperature for 30 minutes, followed by the addition of 10 μL of 6x SDS loading dye. Samples were then boiled at 95°C for 10 minutes to denature the proteins. Hydroxylamine was added to a final concentration of 0.4 M to cleave S-palmitoylation when needed. Proteins were separated by SDS-PAGE, and the fluorescence signal was detected using the ChemiDoc Imaging System (Bio-Rad) with the Rhodamine channel. The gel was subsequently stained with Coomassie Brilliant Blue (B7920, Sigma) to confirm equal protein loading.

### Acyl-biotin exchange (ABE) assay

The ABE protocol was modified as previously described ^53^. Cells were lysed with ABE lysis buffer (100 mM Tris-HCl pH 7.2, 150 mM NaCl, 2.5% SDS, 10 mM N-ethylmaleimide (NEM), protease inhibitors, nuclease) and incubate for 1 hour at room temperature. After quenching with 100 mM 2,3-dimethyl 1,3-butadiene for 1 hour, all the unreacted chemicals were extracted using one-tenth volume of chloroform. The top layer was then divided and mixed with either 0.5 M hydroxylamine or NaCl and treated with Biotin-HPDP. After 1 hour incubation, proteins were precipitated with one volume of chloroform, four volumes of methanol, and three volumes of ddH2O. The resulting protein pellet was washed with 3 volumes of methanol resuspended in ABE resuspension buffer (100 mM Tris-HCl pH 7.2, 150 mM NaCl, 5 mM EDTA, 2.5% SDS, 8 M urea). The resulting protein lysate was then diluted with PBS, and palmitoylated proteins were immunoprecipitated using streptavidin beads. Both immunoprecipitated (IP) and input samples were saved for SDS-PAGE and S-palmitoylated proteins can be detected using western blot.

### Real-time Quantitative PCR

Cellular RNA were extracted using the Total RNA Kit I (Omega Bio-tek) and cDNA was reverse-transcript using the High-Capacity cDNA Reverse Transcription Kit (Thermo Fisher). The qPCR was performed using 2x Universal SYBR Green Fast qPCR Mix (Abclonal) with the QuantStudio™ 7 Flex Real-Time PCR System. Relative gene expression was calculated using the ΔΔCq method and normalized to ACTB.

### Soft agar assay

To measure anchorage-independent cell growth, 2 mL of 0.6% low melting point agarose with cell culture media was added to each well of a 6-well plate as the base layer. Once the agarose had solidified, 5.0 × 10^^3^ NIH 3T3 cells stably expressing either PCDH, KRAS4A G12D, KRAS4A G12D/C180S and KRAS4B G12D were mixed with 0.3% low melting point agarose and plated on top of the solidified 0.6% agarose layer. The new medium was added every 5-7 days. After 14 days of culture, colonies were stained with 200 µL of 1 mg/mL Nitroblue tetrazolium chloride overnight, imaged using ChemiDoc (Biorad) and quantified by ImageJ and CellProfiler.

### 2D cell proliferation assay

Wild-type or DHHC7 knockdown MiaPaca-2, BxPC3, PANC0327 and Tu8988S cells were seeded into a 96-well plate (10000 cells/well). After 48 hours, the cells were treated with CellTiter-Glo® 2.0 regent following the manufacturer’s instructions. To measure the viability of the cells, the luminescence was measured using BioTek Cytation 5 (Agilent).

### Mouse xenograft tumor models

Tumors were established by subcutaneously injecting MiaPaca-2 cells (3 × 10^6 cells per animal), PANC 0327 cells (1 x 10^6 cells per animal), AsPC1 cells (1 x 10^6 cells per animal) and BxPC3 cells (1 x 10^6 cells per animal) into 4-to 6-week-old male NSG mice (Jackson Laboratory, Bar Harbor, US). Control and DHHC7 knockout MiaPaca-2 cells were cultured normally in cell culture for a few passages until the confluency reach >90%. The day before mice injection, all the cells were re-plated on a new plate to ensure maximum viability. To reduce individual variance, control and DHHC7 knockout MiaPaca-2 cells were injected on the left and right abdomen for the same mouse, respectively. Tumor volume (V) was measured every 2-3 days and calculated using the formula V = 1/2 × length × width². At the end of the experiment, mice were dissected and tumors were harvested, imaged, and weighed. All animal procedures adhered to the guidelines of the National Advisory Committee on Laboratory Animal Research and the University’s Institutional Animal Care and Use Committee.

### Electron microscopy (EM)-spatial analysis

Intact apical plasma membrane sheets of baby hamster kidney (BHK) cells expressing GFP-tagged KRAS constructs without/with treatments were attached to copper EM grids. Following fixation with 4% paraformaldehyde (PFA) / 0.1% gluaraldehyde, GFP anchored to the plasma membrane sheets was immunolabeled with anti-GFP antibody coupled to 4.5 nm gold nanoparticles. Gold particles were imaged via transmission EM (TEM) at 100,000x magnification, assigned x / y coordinates in ImageJ within a select 1μm^2^ plasma membrane area. Ripley’s K-function measured the nanoclustering of the gold-labeled GFP-KRAS constructs in the 1μm^2^ plasma membrane area. The goal was to test a null hypothesis: the gold nanoparticles distribute in a random pattern:

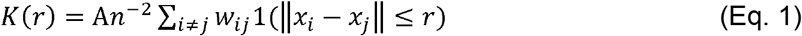

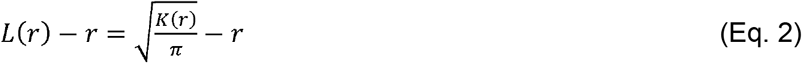

In Equation 1, *K(r*) represents the univariate distribution for a total number of *n* gold particles in a plasma membrane area of *A*; *r* indicates the distance between gold particles with an increment of 1 nm from 1 to 240 nm; ‖ · ‖ represents Euclidean distance, where an indicator of 1(·) is assigned a value of 1 if ‖*x_i_*-*x_j_*‖ ≤ r and a value of 0 if ‖*x_i_*-*x_j_*‖ > r. Edge effects are corrected using a parameter of *w_ij_*^-1^, which indicates the fraction of the circumference of a circle with the center defined as *x_i_* and radius ‖*x_i_*-*x_j_*‖. In Equation 2, *L*(*r*) – *r* represents the linear transformation of *K*(*r*) in Eq. 1. This is accomplished by normalizing *K*(*r*) against the 99% confidence interval (99% C.I.) extracted from Monte Carlo simulations. As signified by the null hypothesis, *L*(*r*) -*r* = 0 when gold particles distribute a complete random pattern. *L*(*r*) -*r >* 99% CI of 1 when gold particles form statistically meaningful clusters. Larger *L*(*r*) -*r* values indicate more extensive clustering. *L_max_* represents the peak values of *L*(*r*) - *r* curves, and are used as a summary statistic. For each condition, at least 15 plasma membrane sheets from individual cells were imaged, analyzed and pooled. Statistical significance was evaluated via comparing our calculated point patterns against 1000 bootstrap samples in bootstrap tests ^54,55^.

### Immunofluorescence and PM fraction calculation

U2OS cells were seeded in 35-mm glass bottom dishes (MatTek), and mScarlett-Lck-N-myristylation together either mNeoGreen-KRAS4A WT or C180S plasmids were overexpressed. After 24 h, cells were fixed with 4% PFA (v/v in PBS) for 15 mins and washed with PBS twice. Samples were then mounted with DAPI Fluoromount-G (0100-20, SouthernBiotech) and imaged using laser scanning inverted confocal microscopy (LSM880, Zeiss). PM fraction was calculated using CellProfiler ^56^ where primary object (nucleus mask) was firstly identified using DAPI channel, and secondary object (whole cell mask) was identified using mScarlett channel. The PM mask was created by shrinking the whole cell mask by 1 pixel size. The PM/Cytosol Ratio was calculated using the average mNeoGreen fluorescence intensity of the corresponding cellular regions.

### Subcellular fraction

Cells were cultured and fractionated following the protocol of Minute™ Plasma Membrane/Protein Isolation and Cell Fractionation Kit (Invent Biotechnologies). The resulting fractions were analyzed using western blot.

### Statistical analysis

Western blot quantifications were obtained from n = 3 independent experiments, and cell proliferation assays were performed with n = 8 replicates per condition; soft agar and qPCR experiments were performed with n = 3. Data are presented as mean ± SD. Statistical significance was determined using a two-tailed Student’s t-test for comparisons between two groups and one-way ANOVA for comparisons among multiple groups; P < 0.05 was considered significant.

## Author Contribution

W.C. contributed to project conception, designed and performed experiments, analyzed data, and wrote the manuscript. S.Y. contributed to computational modeling and manuscript revision. G.M. performed experiments and assisted with experimental design. X.C. contributed to biochemical assays and project design. X.L. and J.Z. assisted with mouse experiments. Y.L. performed qPCR experiments. N.A. and Y.Z. performed the Electron microscopy and data analysis. L.M. and K.F.M. contributed to mouse experiments and data interpretation. M.I.P.M. and M.E.L. performed the in vitro PAT assay. H.L. directed and supervised the study, secured funding, and revised the manuscript. All authors read and approved the manuscript.

## Acknowledgement

We are grateful to Mark Philips for providing the KRAS4A-specific antibody and for helpful comments on the manuscript. This work is supported in part by funding from Howard Hughes Medical Institute, a Dream Team Award from the Cancer Research Foundation and University of Chicago Comprehensive Cancer Center, and Ludwig Center at the University of Chicago.

## Supplementary Figures

**Figure S1.**
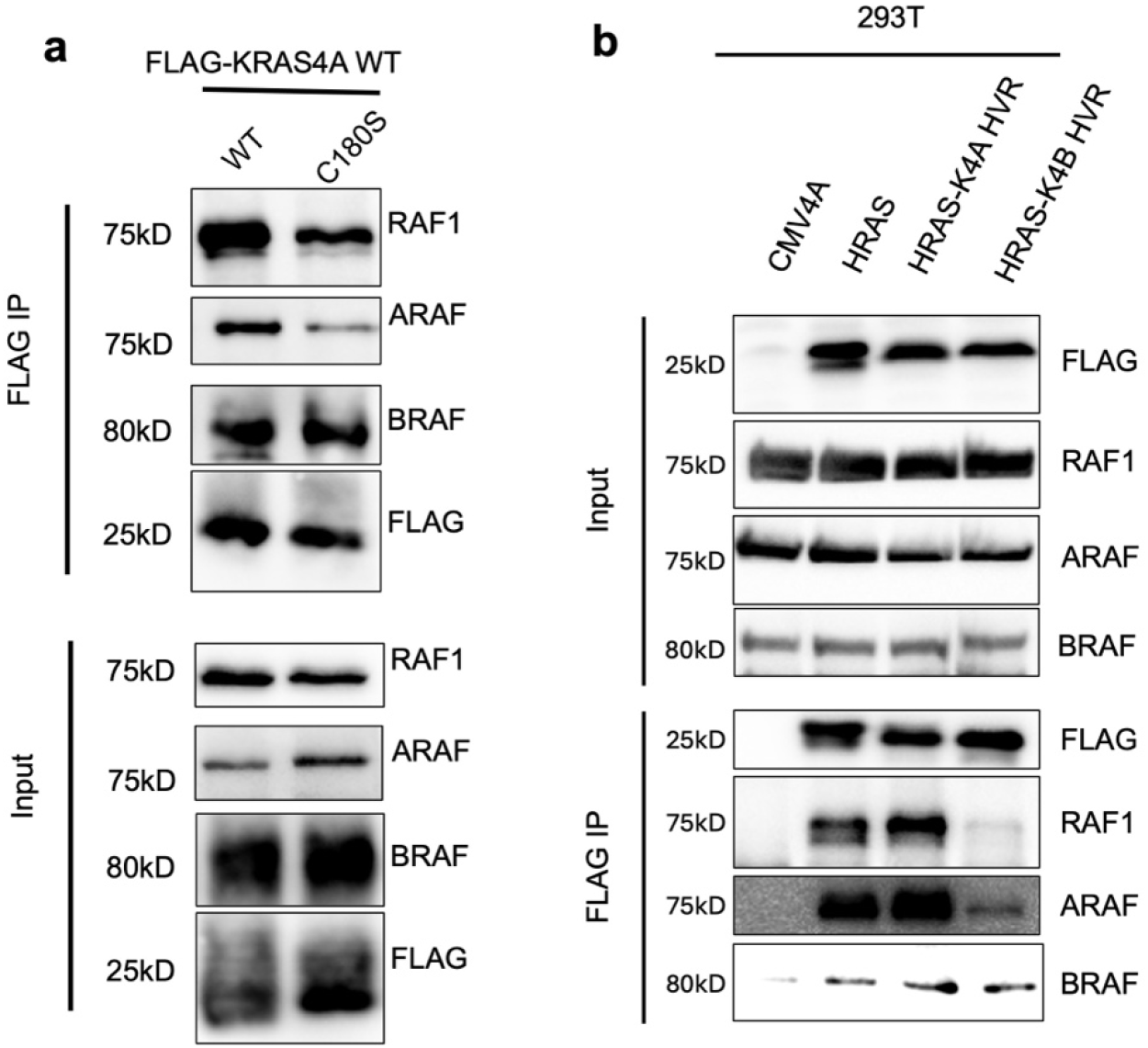
RAF isoform binding to palmitoylation-deficient KRAS4A and to HRAS–KRAS HVR chimeras. (a) Co-immunoprecipitation of RAF1, ARAF, and BRAF with FLAG-KRAS4A WT or C180S expressed in 293T cells. Lysates (input) and FLAG immunoprecipitates were immunoblotted for RAF1, ARAF, BRAF, and FLAG. (b) Co-immunoprecipitation of RAF isoforms with HRAS and with HRAS chimeras carrying the KRAS4A or KRAS4B hypervariable region (HVR). FLAG-tagged constructs were expressed in 293T cells (CMV4A: empty-vector control). Input lysates and FLAG immunoprecipitates were immunoblotted for FLAG, RAF1, ARAF, and BRAF.

**Figure S2.**
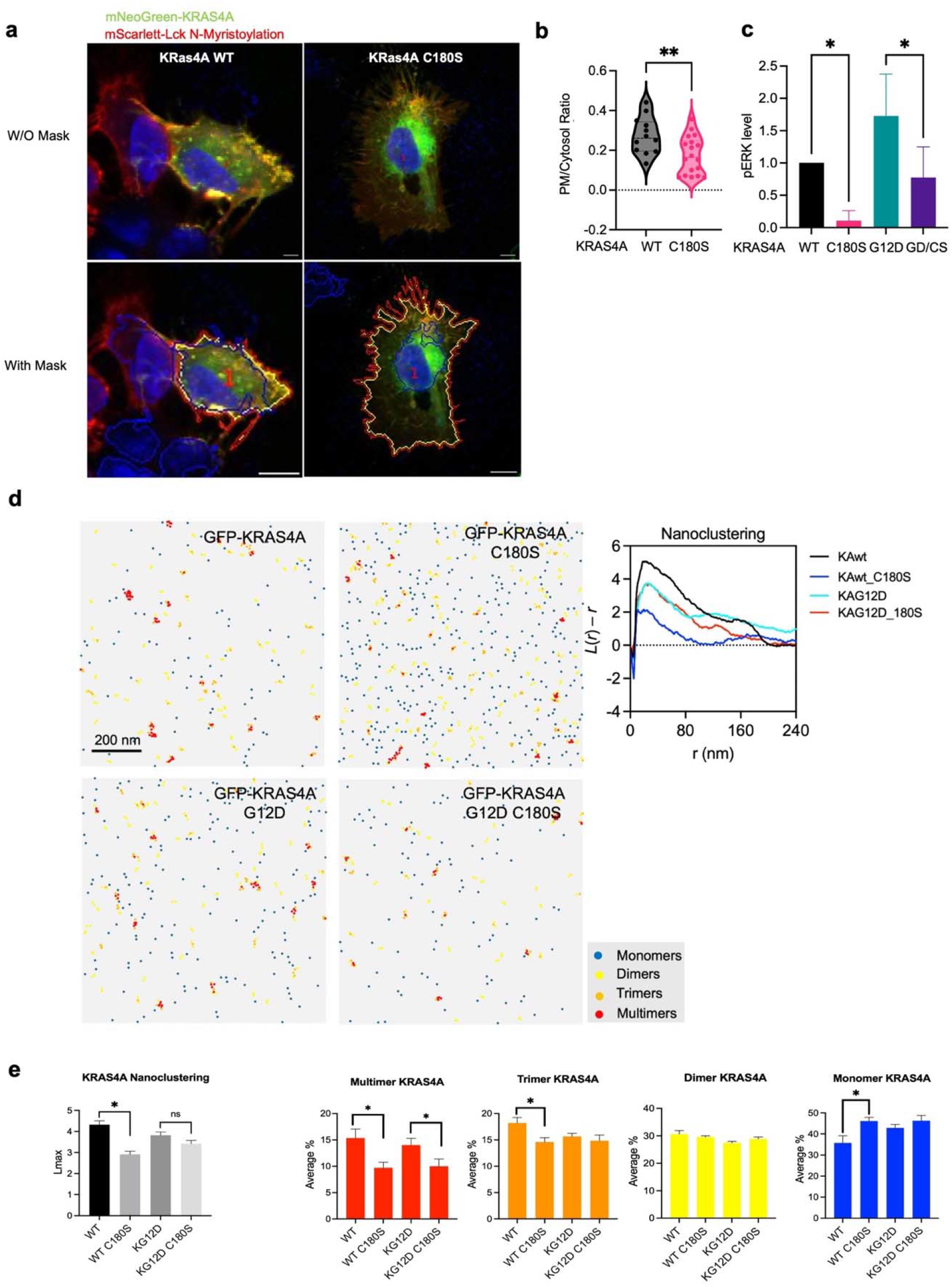
C180 palmitoylation affects KRAS4A plasma membrane localization and nanoclustering. (a) U2OS cells transfected with mNeoGreen-KRAS4A WT, C180S and mScarlett-Lck were imaged using confocal microscope and PM region was identified as the region between whole cell mask (red) and cytosol mask (yellow) mask. (b) Plasma membrane/cytosol ratio of KRas4A was quantified using mean fluorescence intensity of different cellular compartments. (c) The quantification of pERK level difference in Figure 1c. (d) The nanoclustering of KRAS4A was measured using Immunolabeled PM sheet under electron microscopy and quantified using in-house software. Representative images of immunogold-labeled plasma membrane sheets are shown for GFP-KRAS4A WT, C180S, G12D, and G12D/C180S, with gold particles classified as monomers, dimers, trimers, or multimers (scale bar, 200 nm), together with the L(r) − r analysis for each condition. The overall nanoclustering quantified and individual oligomer was quantified in (e). The results in (b), (c), (d) and (e) are shown as mean ± SD. ns, not significant. *P < 0.05; **P < 0.01; ***P < 0.001. Western blot quantification in (c) is from n = 3 independent experiments. Statistical significance was determined by two-tailed Student’s t-test for comparisons between two groups and by one-way ANOVA for comparisons among multiple groups.

**Figure S3.**
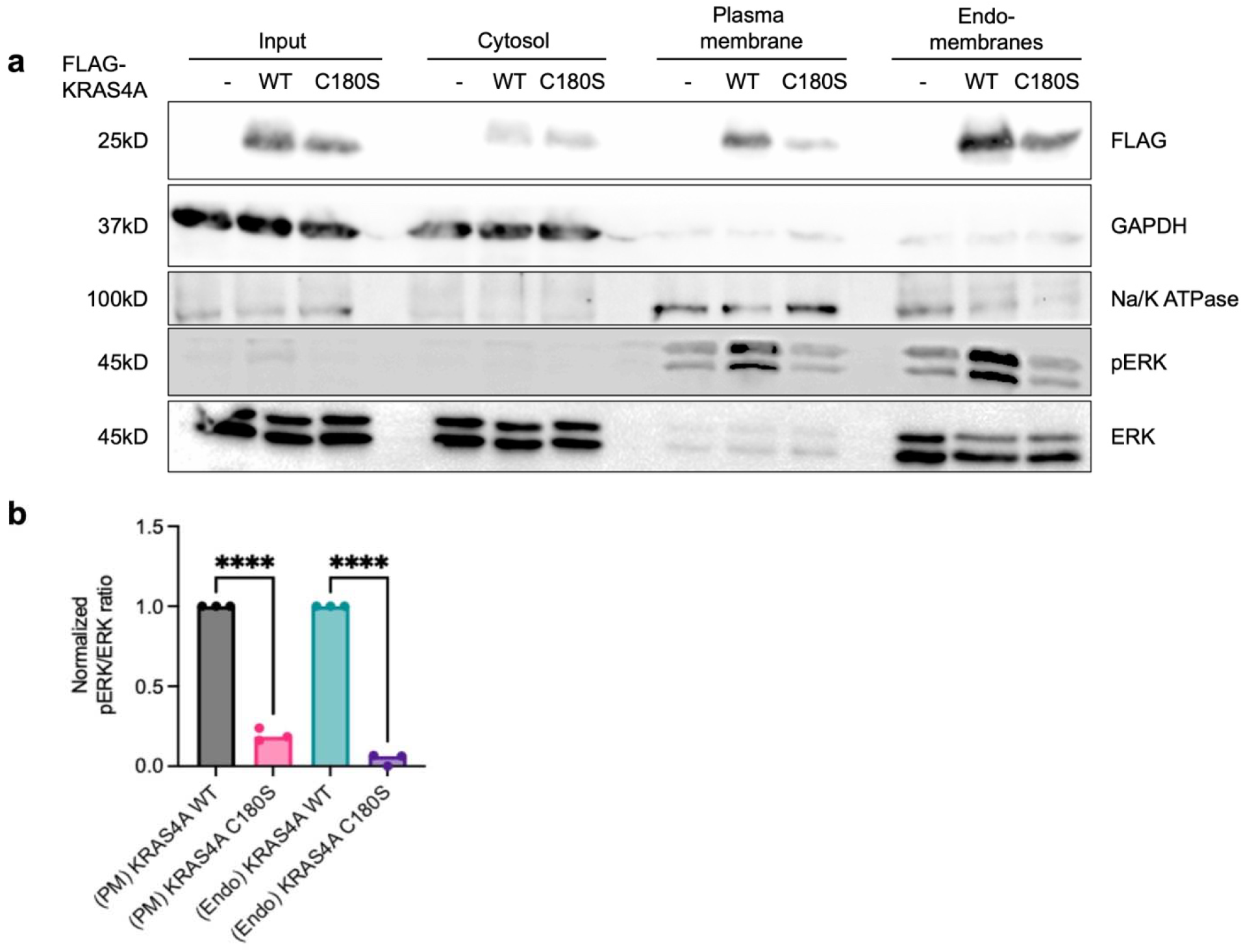
The membrane-localized KRAS4A C180S mutant is functionally impaired. Cells expressing FLAG-KRAS4A WT or C180S (– indicates no KRAS4A) were separated into cytosol, plasma membrane, and endomembrane fractions and immunoblotted for FLAG, GAPDH (cytosol marker), Na/K ATPase (plasma membrane marker), pERK, and ERK. Bottom: quantification of the pERK/ERK ratio in the plasma membrane (PM) and endomembrane (Endo) fractions, normalized to the input level of FLAG-KRAS4A WT or C180S and shown relative to WT in each fraction (n = 3; one-way ANOVA, ****P < 0.0001).

**Figure S4.**
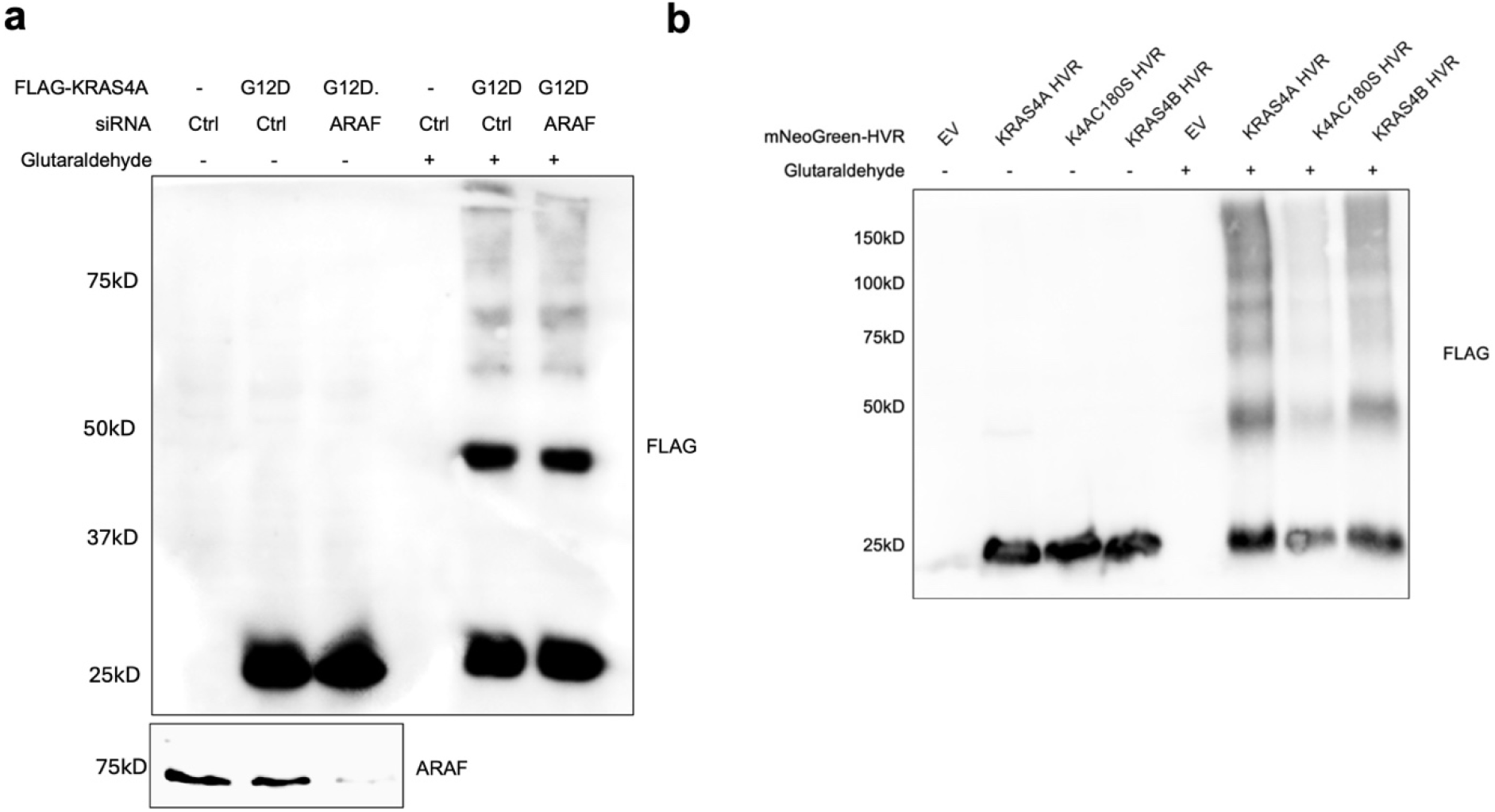
Controls for KRAS4A nanoclustering. (a) ARAF knockdown does not reduce KRAS4A crosslinking. FLAG-KRAS4A G12D was expressed with control or ARAF siRNA, treated with or without glutaraldehyde, and immunoblotted for FLAG and ARAF. (b) Glutaraldehyde crosslinking of the HVR of KRAS4A WT, KRAS4A C180S, or KRAS4B fused to mNeonGreen. Cells expressing empty vector (EV) or the indicated FLAG-tagged mNeonGreen-HVR fusions were treated with or without glutaraldehyde and immunoblotted for FLAG.

**Figure S5.**
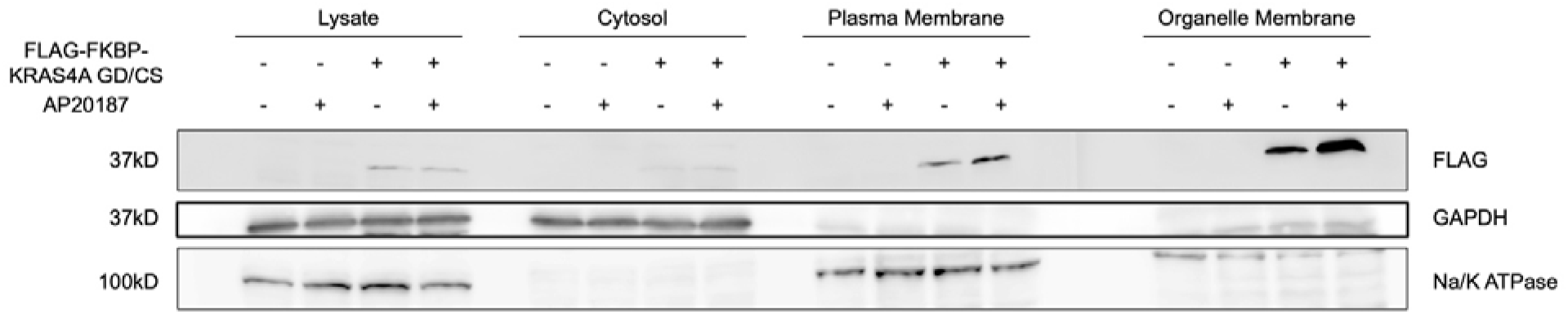
Subcellular fractionation of cells expressing FLAG-FKBP-KRAS4A GD/CS with and without AP20187. The FLAG-FKBP-KRAS4A GD/CS levels in total lysate, cytosolic, plasma membrane, and organelle membrane fractions detected by western blot for FLAG and different markers.

**Figure S6.**
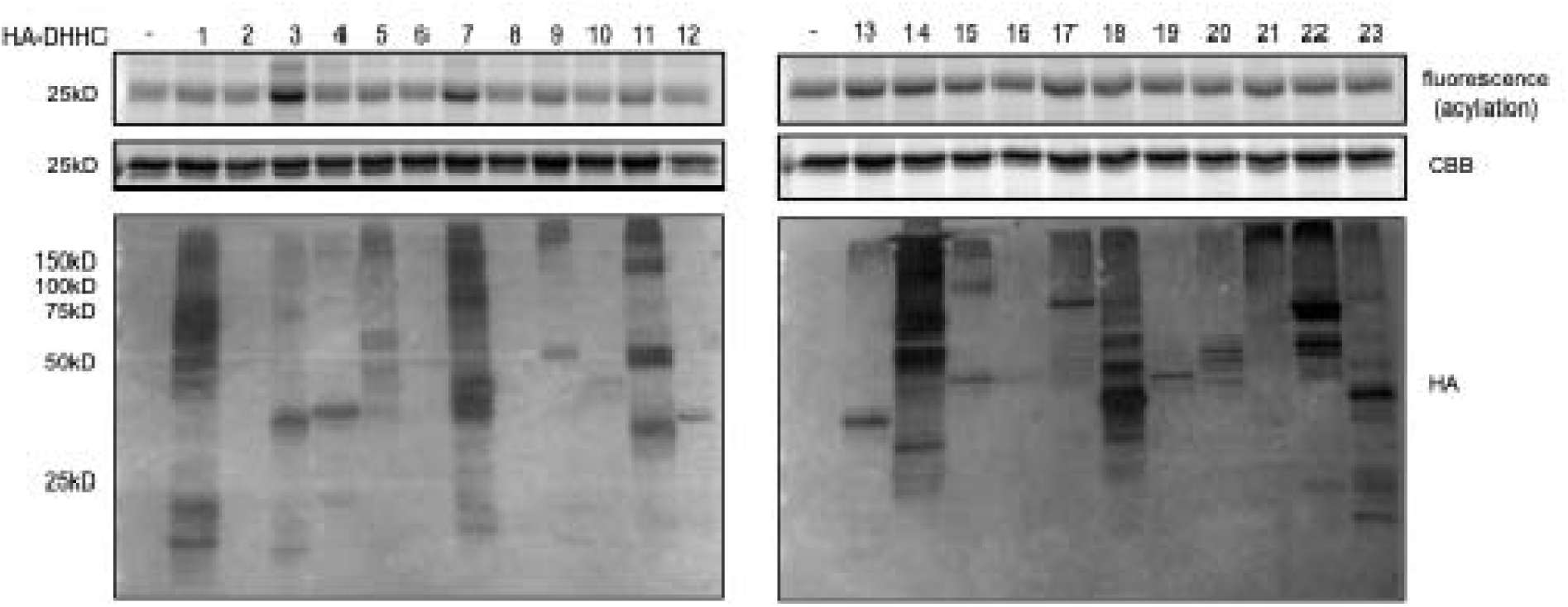
Overexpression of DHHC7 and DHHC3 promotes KRAS4A palmitoylation. HEK293T cells were transfected with FLAG-KRAS4A and 23 HA-DHHCs. The S-palmitoylation level was detected using Alkyne-14 metabolic labeling.

**Figure S7.**
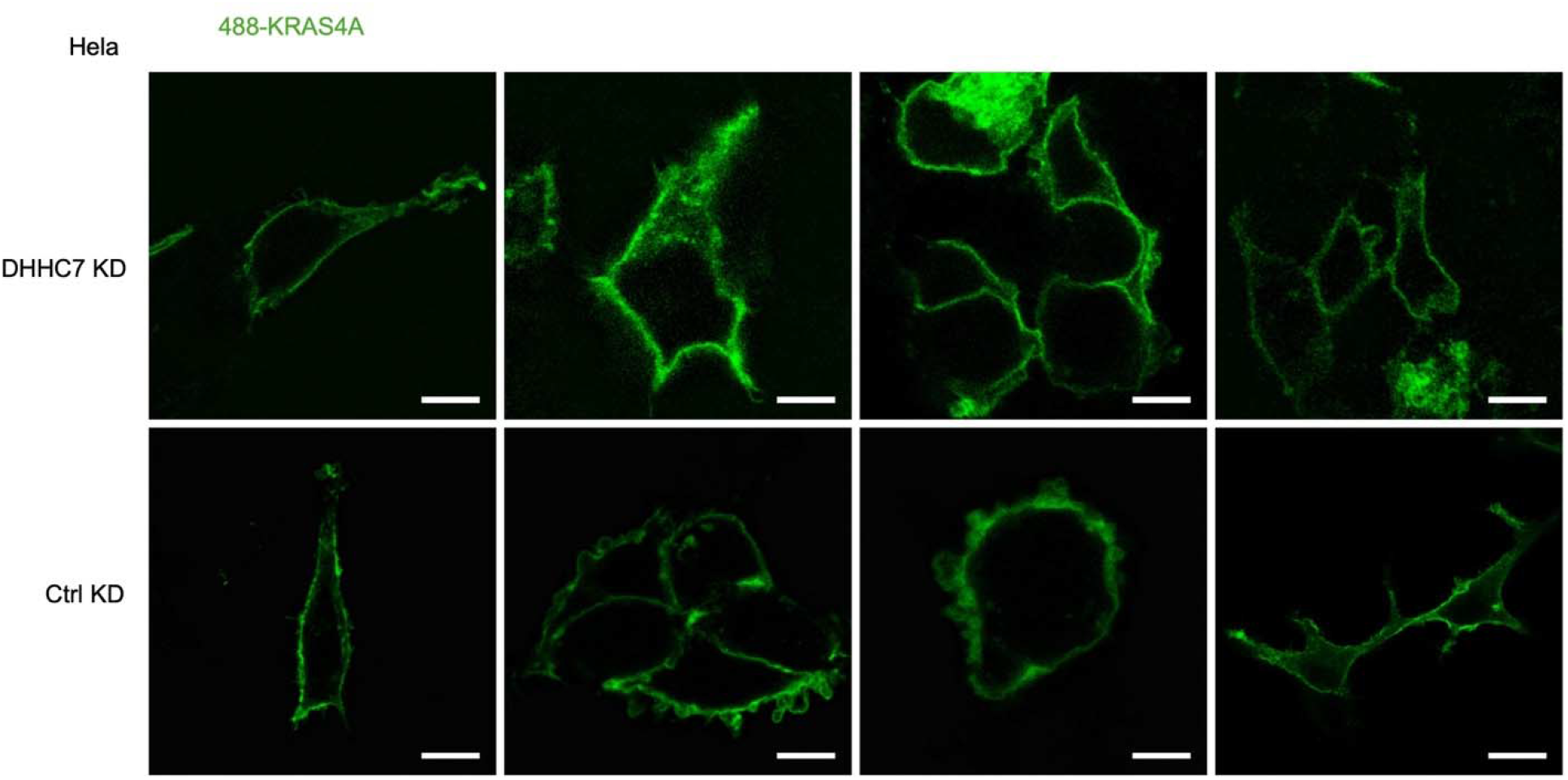
Confocal images of KRAS4A in HeLa cells after DHHC7 knockdown (KD) or control (Ctrl) knockdown. Scale bar, 10 µm.

**Figure S8.**
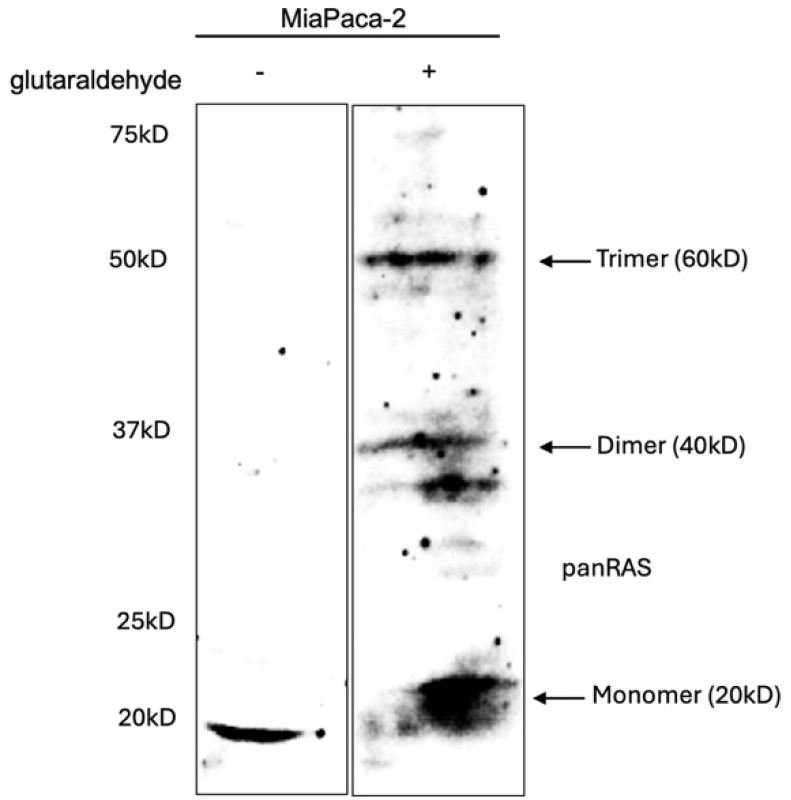
Crosslinking of endogenous KRAS. Glutaraldehyde crosslinking of endogenous KRAS in ZDHHC7 WT and KO MiaPaca-2 cells, detected with a pan-KRAS antibody.

**Figure S9.**
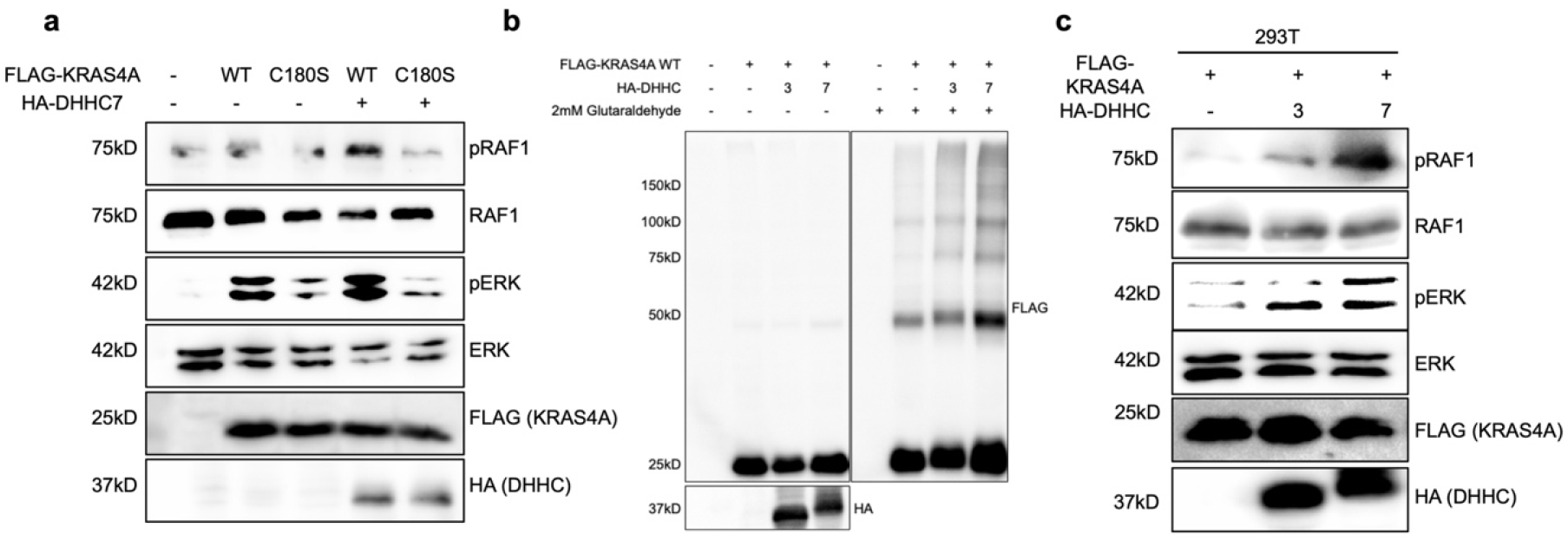
DHHC7 overexpression increases pRAF1, pERK, and KRAS4A oligomerization. (a) DHHC7 overexpression enhances pRAF1 and pERK signaling in cells expressing KRAS4A WT but not C180S. (b) Overexpression of DHHC3 and DHHC7 promotes KRAS4A nanoclustering, detected by glutaraldehyde crosslinking. (c) 293T cells expressing FLAG-KRAS4A were co-transfected with empty vector (-), HA-DHHC3 (3), or HA-DHHC7 (7) and immunoblotted for pRAF1, RAF1, pERK, ERK, FLAG, and HA.

**Figure S10.**
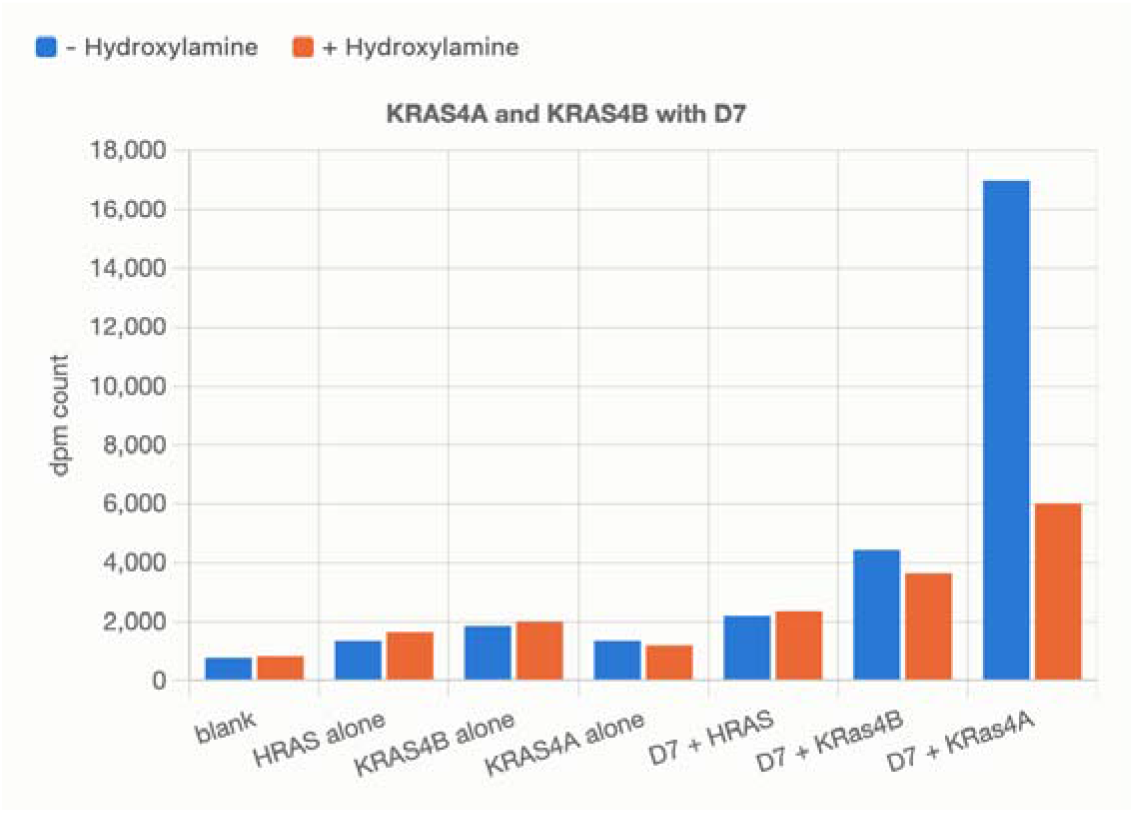
In vitro palmitoylation (PAT) assay with purified DHHC7 and KRAS4A, KRAS4B, or HRAS. Purified DHHC7 (50 nM final) was incubated with 2 µM purified substrate (KRAS4A, KRAS4B, or HRAS) and 1 µM [^3^H]palmitoyl-CoA in a 50 µL reaction at room temperature for 10 min, and the reaction was quenched in SDS loading dye containing 25 mM TCEP. Samples were resolved by SDS-PAGE, the Coomassie-stained substrate bands were excised, dissolved in Soluene, and ^3^H incorporation was quantified by liquid scintillation counting; values are reported as disintegrations per minute (dpm) after subtracting the background measured for substrate incubated without enzyme, which controls for non-enzymatic acylation. Each reaction was split and analyzed without (-) or with (+) hydroxylamine (HA) treatment. Because HA cleaves thioester bonds but not amide bonds, the HA-sensitive fraction (the difference between -HA and + HA) is the readout for genuine cysteine S-palmitoylation, whereas residual + HA signal reflects HA-resistant (e.g., amide-linked) or nonspecific label incorporation.

**Figure S11.**
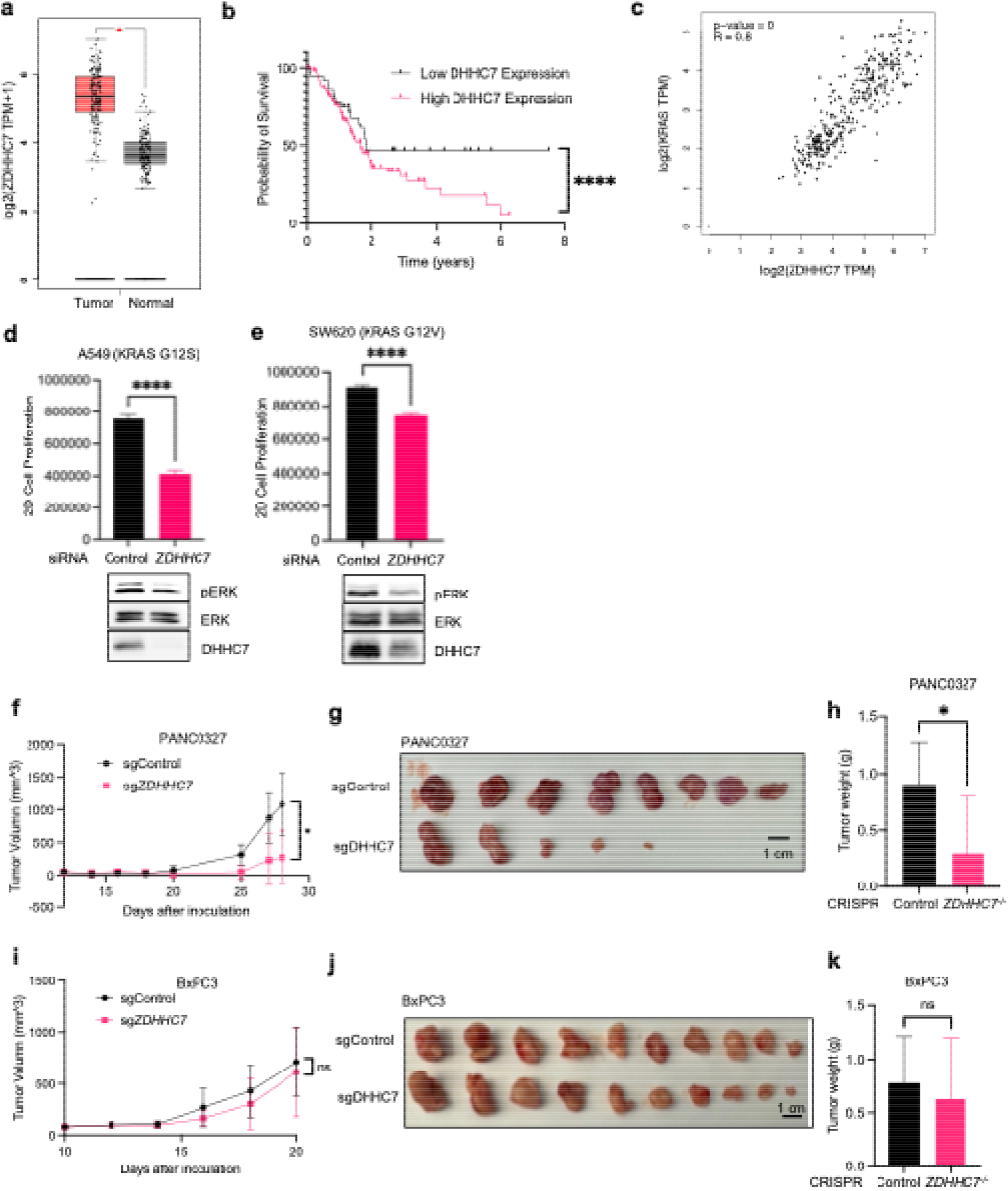
DHHC7 knockout decrease tumor proliferation in PANC0327 but not in BxPC3 xenograft. (a) The pancreatic expression of *ZDHHC7* between TCGA PAAD normal (n=179) and tumor (n=171) tissues were analyzed and plot as box plot with median, upper quartile, lower quartile and 95% confidence interval using the GEPIA web tool. (b) Kaplan–Meier survival curve of 176 pancreatic cancer patients in TCGA databases analyzed using the Human Protein Atlas. Patients were divided into two groups (cut off = 20.53) based on *ZDHHC7* mRNA levels in their tumors. (c) The gene correlation plot between *ZDHHC7* and *KRAS* was analyzed using TCGA PAAD patients (n=350) and GTEx pancreas dataset using GEPIA web tool. The Pearson coefficient was calculated. *ZDHHC7* was knockdown using siRNA in two KRAS-driven cancer cell lines (d) A549 and (e) SW620. 2D cell growth was measured using Cell Titer Glow after 48-72 hrs. (f) Pancreatic cancer xerograph was performed using genetically modified PANC0327 cell line with Control/*ZDHHC7* CRISPR knockout. The tumor volume was measured and quantified every 2-4 days. (g) The mice were dissected after 30 days and (h) final tumors were weighted. (i) Pancreatic cancer xerograph was performed using genetically modified BxPC3 cell line with Control/*ZDHHC7* CRISPR knockout. The tumor volume was measured and quantified every 2-4 days. (j) The mice were dissected after 30 days and (k) final tumors were weighted. The results in (a), (d), (e), (f), (h), (i) and (k) are shown as mean ± SD. ns, not significant. *P < 0.05; **P < 0.01; ***P < 0.001. Cell proliferation in (d) and (e): n = 8. Statistical significance was determined by two-tailed Student’s t-test for comparisons between two groups and by one-way ANOVA for comparisons among multiple groups.

**Figure S12.**
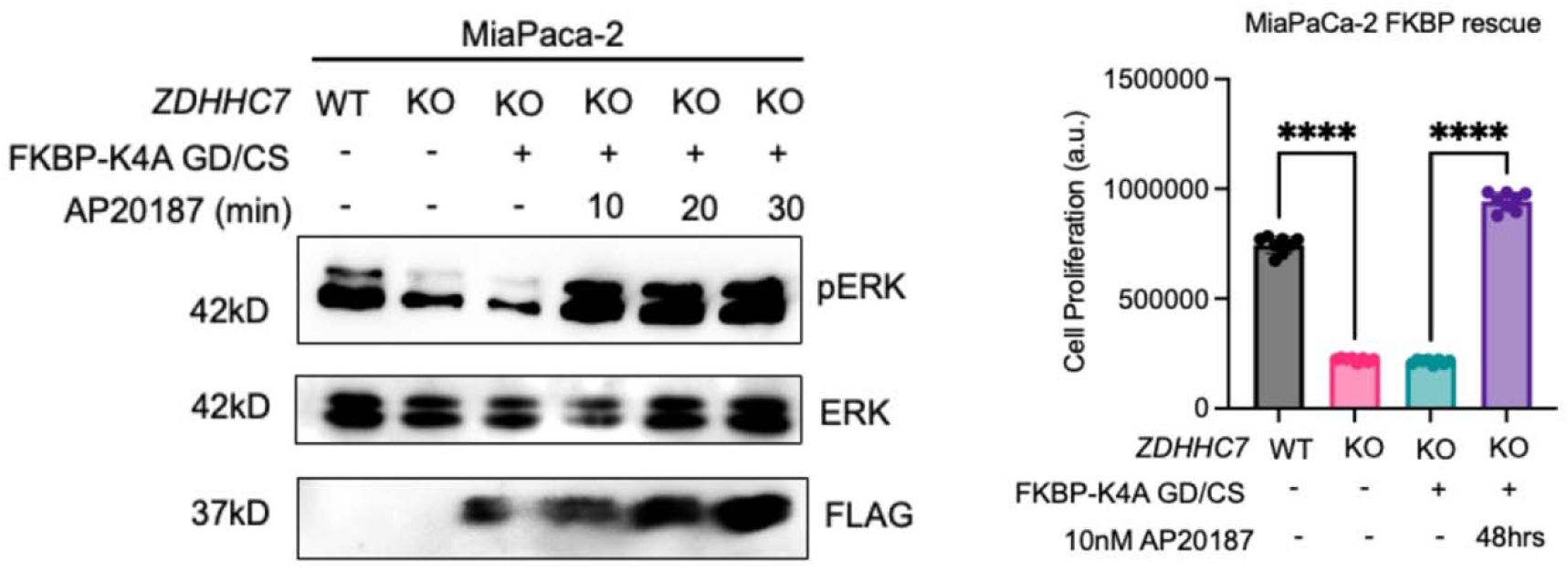
Clustering of FKBP-KRAS4A G12D/C180S rescues ZDHHC7 knockout in MiaPaca-2 cells. Left: pERK immunoblot of *ZDHHC7* WT and KO MiaPaca-2 cells expressing FKBP-KRAS4A G12D/C180S, treated with AP20187 for the indicated times. Right: proliferation of the same cells after 48 hours with or without 10 nM AP20187 (n = 8; one-way ANOVA, ****P < 0.0001). The immunoblot is from a single experiment.

## Supplementary Note 1. Thermodynamic model linking KRAS4A localization and oligomerization

## Background: membrane localization driving self-assembly is an established protein behavior

The mechanism set out below is not specific to KRAS4A, and it is not a new proposal. That membrane localization drives the assembly of otherwise soluble proteins is an established behavior of membrane-targeting proteins, and it has been derived from equilibrium thermodynamics and validated quantitatively against measured data by Johnson (PLoS Comput Biol 2018, 14:e1006031; PMID: 29505559). The binding scheme analyzed in that work is reproduced in Figure SN1.

**Figure SN1.**
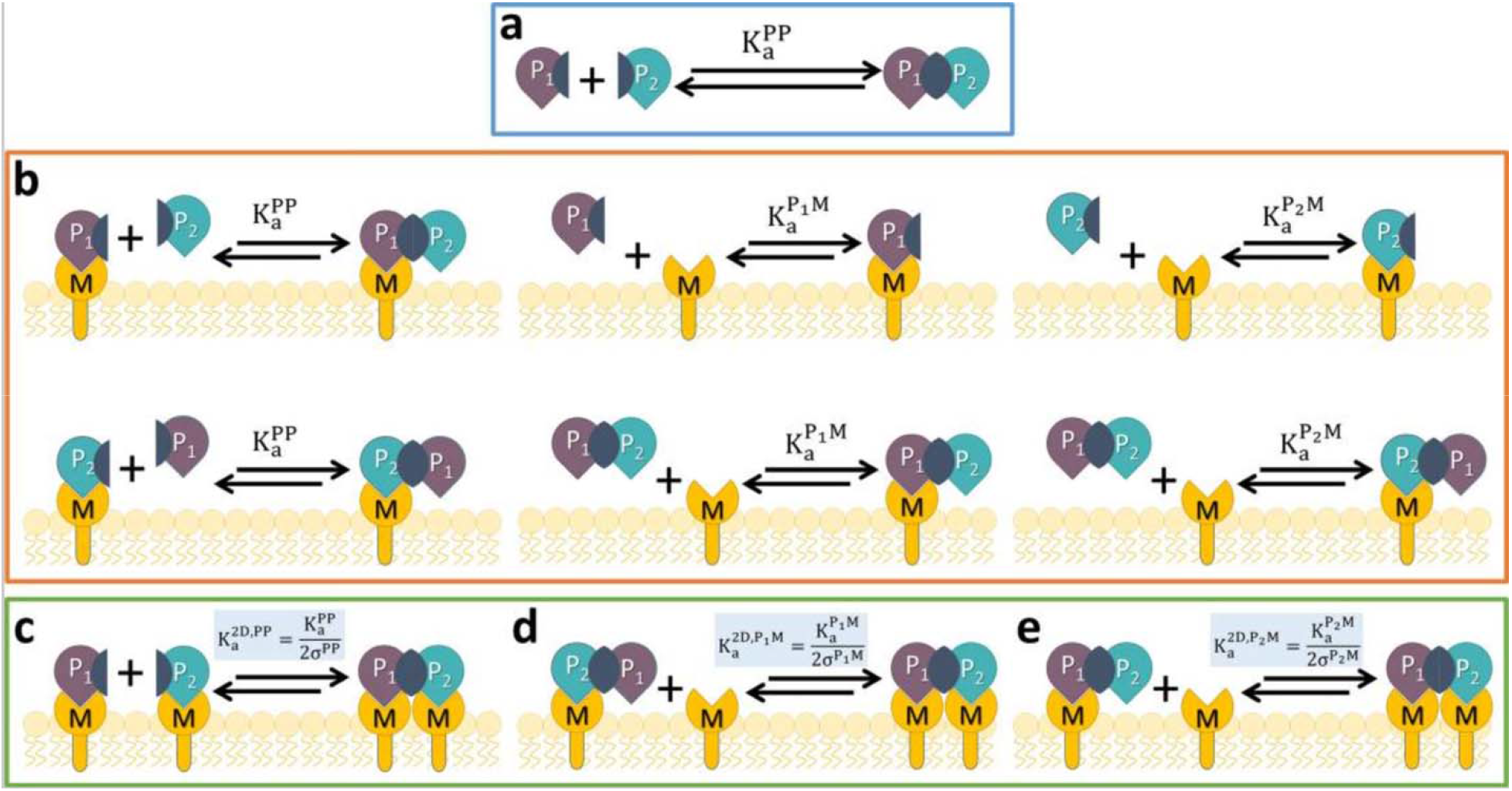
The binding scheme underlying the established behavior, reproduced from Johnson, PLoS Comput Biol 2018, 14:e1006031 (Figure 1; CC BY 4.0). Two soluble proteins, P1 and P2, bind each other and also bind a membrane lipid, M. (a) Protein–protein binding in solution, governed by the three-dimensional constant Ka^PP^. (b) Reactions that draw protein to the surface: solution-phase binding to an already membrane-bound partner, and lipid binding b free protein or by a preformed P1P2 complex. (c–e) Reactions between species that are already on the membrane and therefore proceed in two dimensions, with Ka^2D^ = Ka^3D^/(2σ) for each pair. Panels (d) and (e) are the cooperative arm of the cycle: a protein–protein complex engages a second lipid, so complexation itself stabilizes membrane residence. For KRAS4A the same topology applies with the two lipid anchors, farnesyl and the C180 palmitoyl group, in place of M.

This work establishes the behavior in both directions. In one direction, confining two binding partners to a bilayer reduces the dimensionality of their search and so raises their effective affinity; two-dimensional and three-dimensional equilibrium constants are related by Ka^2D^ = Ka^3D^/(2σ), where σ is a length absorbing the change in standard state and the largely entropic cost of surface restriction. Using measured protein–protein and protein–lipid affinities, cellular protein and lipid concentrations, and volume-to-surface-area ratios for 46 interactions among 37 membrane-targeting proteins, that study found effective-affinity enhancements of 10–1000-fold upon localization, with the magnitude set by lipid concentration, the volume-to-surface-area ratio, and the 3D-to-2D conversion. In the reverse direction, the same analysis shows the number of molecules retained on the membrane depends on the protein–protein interaction strength: a species engaging the bilayer through more than one contact is held better than its separate subunits. The two directions close into a positive loop, and the loop is observed experimentally: BAR-domain proteins such as endophilin and FCHo1/2 oligomerize appreciably only once on the membrane, and homodimer-forming partners build larger oligomers that feed back into stronger membrane stabilization.

KRAS4A nanoclustering fits this framework directly. KRAS4A is a small protein that partitions between cytosol and plasma membrane and self-associates into nanoclusters on the membrane, so it is exactly the class of protein the analysis above describes. Palmitoylation at C180 supplies a second lipid anchor and increases the membrane-bound fraction, which by the first direction should raise self-association; the nanoclusters so formed carry two anchors per subunit and, by the second direction, should in turn be retained on the membrane better than monomers. Our observations follow that expectation: palmitoylated KRAS4A shows increased crosslinked oligomer and nanoclustering by EM relative to the C180S mutant, which has lower membrane binding, and the signaling difference tracks the clustering difference rather than palmitoylation itself, and as such, enforced dimerization of the FKBP-tagged GD/CS mutant restores pERK without restoring the modification. The magnitudes are also compatible: the 10–1000-fold enhancement in effective affinity reported for localization is more than sufficient to account for the clustering and MAPK differences we observe.

The remainder of this note makes that argument explicit for KRAS4A, as a closed thermodynamic cycle relating the four species involved.

### 1. The two processes sit on one cycle

KRAS4A can be a monomer (*M*) or a dimer (*D*), and either cytosolic (*c*) or membrane-bound (*m*). This gives four species and four equilibria: dimerization in the cytosol (*K_1_*) and on the membrane (*K_1_*′), and localization of the monomer (*K_2_*) and of the dimer (*K_2_*′). Membrane species are counted per area, cytosolic ones per volume.

**Figure SN2.**
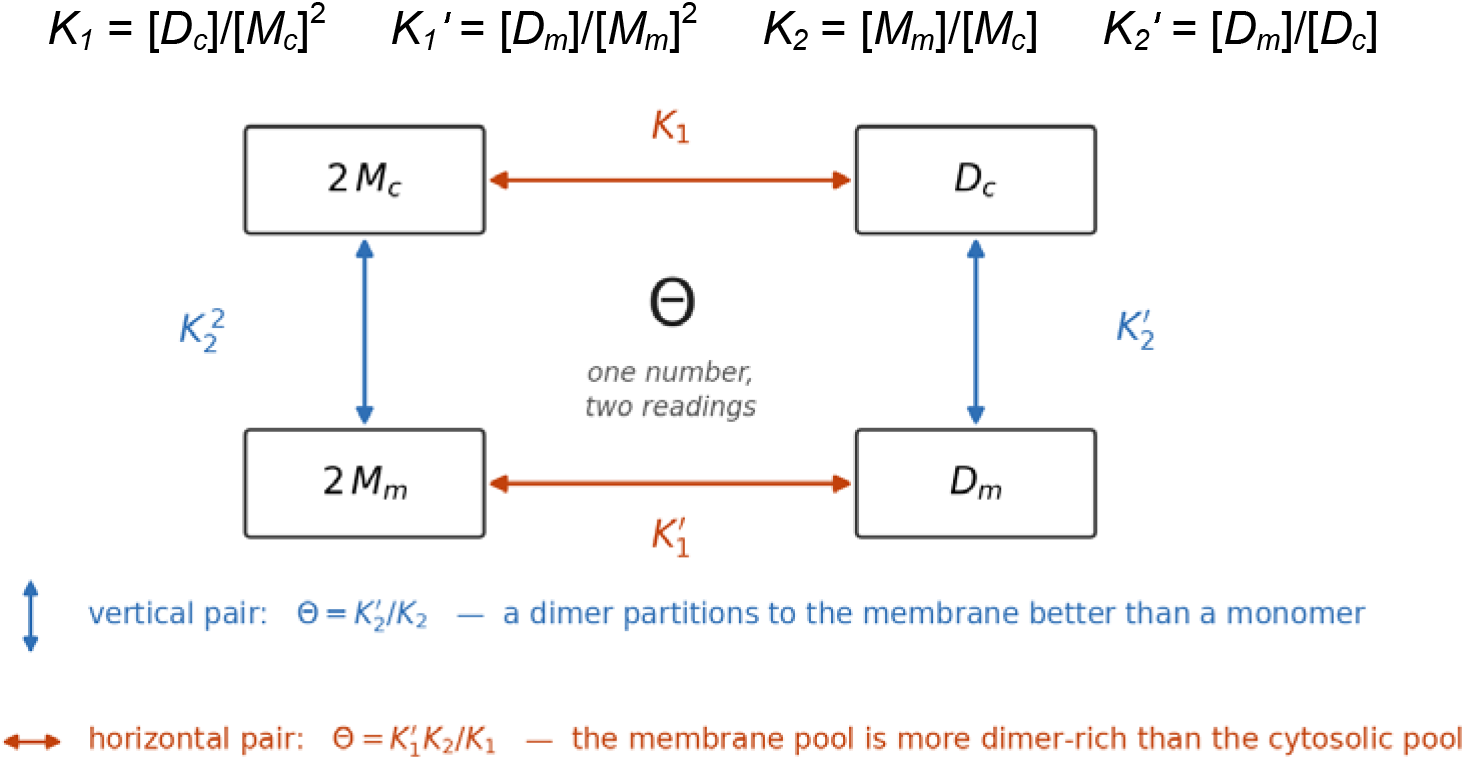
The four species form a closed cycle. Comparing the vertical arrows asks about localization; comparing the horizontal arrows asks about dimerization. Note the stoichiometry: K_2_ is defined per monomer, so the left vertical arrow, which transfers two monomers, carries K_2_², whereas the right vertical arrow transfers a single dimer and carries K_2_′. Θ = K_2_′/K_2_ compares one dimer with one monomer and therefore contains no square.

### 2. The cycle ties the two processes together

*D_m_* is a single state, so the two routes from 2*M_c_* — dimerize then localize, or localize then dimerize — must give the same overall constant:

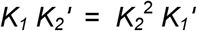

This only assumes that the system is at equilibrium. Rearranged, it defines a single quantity:

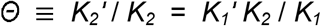

The left form (*K_2_*′ / *K_2_*) **asks how much better a dimer sticks to the membrane than a monomer**. The right form, after substituting *K_2_* = [*M_m_*]/[*M_c_*], is ( [*D_m_*]/[*M_m_*]) / ( [*D_c_*]/[*M_c_*]), **asks how much more of the membrane pool is dimeric than of the cytosolic pool.**

**Localization and oligomerization therefore cannot be changed independently: whatever alters one also alters the other, because of this equation.**

### 3. A dimer binds the membrane better than a monomer

A dimer binds the membrane better than a monomer for two reasons. First, it carries two anchors and can remain membrane-bound when either one is inserted, whereas a monomer is released as soon as its single anchor leaves the bilayer. Second, the two insertions are not independent: once the first anchor is inserted, the partner subunit is tethered alongside it and its own anchor already sits within reach of the bilayer, so it no longer has to find the membrane from the cytosol. This is the familiar avidity effect of multivalent membrane binding: the second anchor inserts more readily than the single anchor of a free monomer.

Both effects act in the same direction, so a dimer partitions to the membrane more strongly than a monomer: Θ >.

**Figure SN3.**
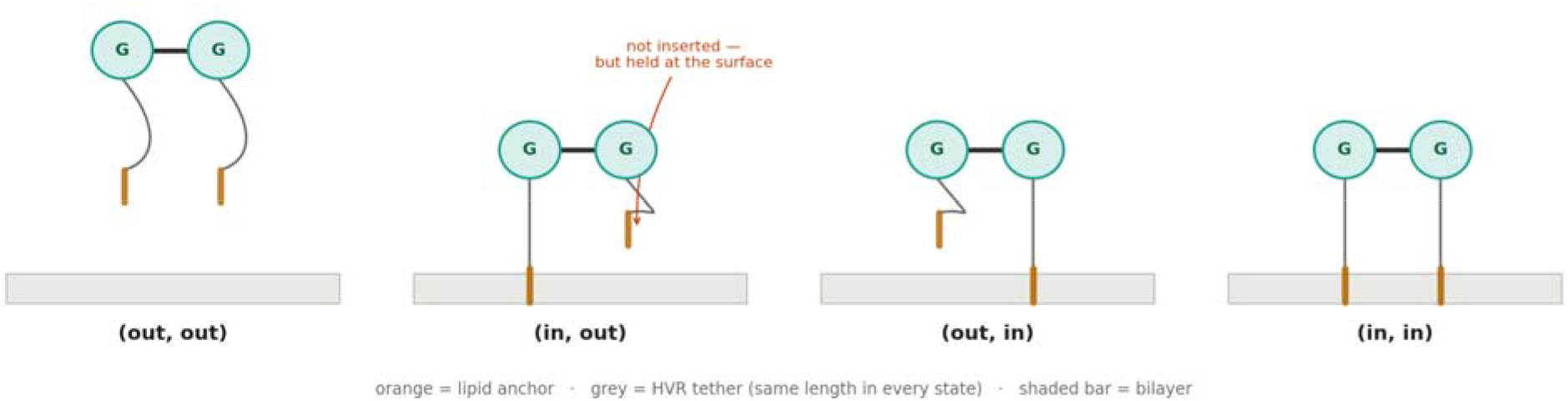
Of the four configurations available to a dimer, three keep it on the membrane; a monomer has only one way to hold on.

### 4. Therefore localization drives oligomerization

Because Θ is one number, Θ > 1 read on the other side of the cycle says

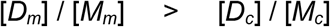

#### KRAS4a on the membrane is more dimeric than KRAS4a in the cytosol

Localization does not merely accompany dimerization; it makes dimerization more favorable, and by exactly the factor by which two anchors beat one.

### 5. Localization and dimerization form a closed loop

The KRAS4A dimer carries more membrane anchors, so it holds the membrane more tightly than the monomers. Localization promotes dimerization, and the dimerization it produces strengthens localization. So membrane localization and dimerization form a closed loop and strengthen each other.

### 6. Palmitoylation is an entry into the loop

Palmitoylation supplies the second lipid anchor of KRAS4a and enhances binding to the plasma membrane; in terms of the cycle, it raises *K_2_*. Because *K_2_* appears in Θ = *K_1_*′*K_2_*/*K_1_*, raising it raises the dimer content of the membrane pool; and the dimers so formed carry more membrane anchor, so they localize to membrane better still. **Palmitoylation therefore does not act on localization alone, it also increases dimerization (and higher orde oligomerization).** A palmitoylation-deficient protein is retained on the membrane by its farnesyl anchor alone, so *K_2_* is small and the loop starts only weakly.

### 7. Experimental Evidence

This model makes the rescue experiment a direct test rather than an anomaly. If palmitoylation acts by raising *K_2_* increasing the membrane-bound fraction, it acts as an input to the loop rather than as a chemical requirement for signaling. Enforced dimerization bypasses that input: AP20187 crosslinking of the FKBP-tagged GD/CS mutant produces dimers directly, without restoring the lipid modification that would ordinarily generate them. We observed that pERK returns to the level of the palmitoylation-competent GD construct, indicating that enforced dimerization can substitute for the palmitoyl group in placing KRAS4A within the amplifying loop of localization and oligomerization. Subcellular fractionation of cells expressing FLAG-FKBP-KRAS4A GD/CS (**Figure SN4**) points in the same direction at the level of localization: AP20187 appears to increase its amount in the plasma membrane and organelle membrane fractions, while total FLAG in the lysate is unchanged and the cytosolic signal remains faint. This is what the cycle predicts, because enforced dimers should bind the membrane better than monomers (*K_2_*′ > *K_2_*). The FLAG-FKBP-KRAS4A GD/CS mutant has membrane binding affinity even without AP20187, presumably through its farnesyl anchor, and dimerization by AP20187 further enhances membrane binding. Palmitoylation is thus positioned as a physiological entry point to the loop and enforces dimerization/oligomerization to promote KRAS4A signaling.

**Figure SN4.**
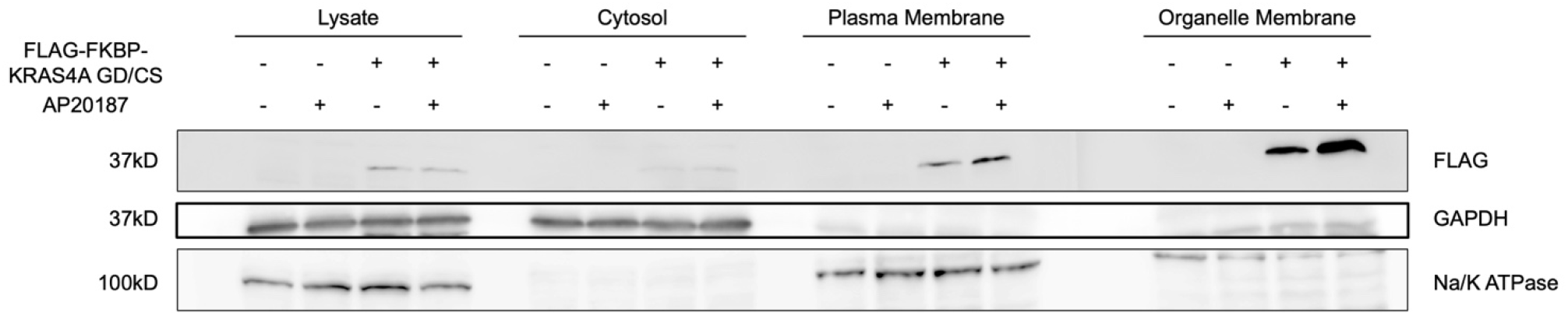
Subcellular fractionation of cells expressing FLAG-FKBP-KRAS4A GD/CS with and without AP20187. The FLAG-FKBP-KRAS4A GD/CS levels in total lysate, cytosolic, plasma membrane, and organelle membrane fractions detected by western blot for FLAG and different markers.

## References

1. Fernández-Medarde, A. & Santos, E. Ras in cancer and developmental diseases. Genes Cancer 2, 344–358 (2011).

2. Simanshu, D. K., Nissley, D. V. & McCormick, F. RAS proteins and their regulators in human disease. Cell 170, 17–33 (2017).

3. Spencer-Smith, R. & O’Bryan, J. P. Direct inhibition of RAS: Quest for the Holy Grail? Semin. Cancer Biol. 54, 138–148 (2019).

4. Isermann, T., Sers, C., Der, C. J. & Papke, B. KRAS inhibitors: resistance drivers and combinatorial strategies. Trends Cancer 11, 91–116 (2025).

5. Drizyte-Miller, K., Talabi, T., Somasundaram, A., Cox, A. D. & Der, C. J. KRAS: the Achilles’ heel of pancreas cancer biology. J. Clin. Invest. 135, (2025).

6. Whitley, M. J. et al. Comparative analysis of KRAS4a and KRAS4b splice variants reveals distinctive structural and functional properties. Sci. Adv. 10, eadj4137 (2024).

7. Tsai, F. D. et al. K-Ras4A splice variant is widely expressed in cancer and uses a hybrid membrane-targeting motif. Proc Natl Acad Sci USA 112, 779–784 (2015).

8. Zhang, X., Cao, J., Miller, S. P., Jing, H. & Lin, H. Comparative Nucleotide-Dependent Interactome Analysis Reveals Shared and Differential Properties of KRas4a and KRas4b. ACS Cent. Sci. 4, 71–80 (2018).

9. Zhao, H. et al. Roles of palmitoylation and the KIKK membrane-targeting motif in leukemogenesis by oncogenic KRAS4A. J. Hematol. Oncol. 8, 132 (2015).

10. Laude, A. J. & Prior, I. A. Palmitoylation and localisation of RAS isoforms are modulated by the hypervariable linker domain. J. Cell Sci. 121, 421–427 (2008).

11. Terrell, E. M. et al. Distinct Binding Preferences between Ras and Raf Family Members and the Impact on Oncogenic Ras Signaling. Mol. Cell 76, 872–884.e5 (2019).

12. Das, T., Yount, J. S. & Hang, H. C. Protein S-palmitoylation in immunity. Open Biol. 11, 200411 (2021).

13. Lin, H. Protein cysteine palmitoylation in immunity and inflammation. FEBS J. 288, 7043–7059 (2021).

14. Simanshu, D. K., Philips, M. R. & Hancock, J. F. Consensus on the RAS dimerization hypothesis: Strong evidence for lipid-mediated clustering but not for G-domain-mediated interactions. Mol. Cell 83, 1210–1215 (2023).

15. Prior, I. A., Muncke, C., Parton, R. G. & Hancock, J. F. Direct visualization of Ras proteins in spatially distinct cell surface microdomains. J. Cell Biol. 160, 165–170 (2003).

16. Li, W. et al. Plasma membrane-associated ARAF condensates fuel RAS-related cancer drug resistance. Nat. Chem. Biol. 21, 1226–1237 (2025).

17. Spencer, D. M., Wandless, T. J., Schreiber, S. L. & Crabtree, G. R. Controlling signal transduction with synthetic ligands. Science 262, 1019–1024 (1993).

18. Nan, X. et al. Ras-GTP dimers activate the Mitogen-Activated Protein Kinase (MAPK) pathway. Proc Natl Acad Sci USA 112, 7996–8001 (2015).

19. S Mesquita, F., et al. Mechanisms and functions of protein S-acylation. Nat. Rev. Mol. Cell Biol. 25, 488–509 (2024).

20. Swarthout, J. T. et al. DHHC9 and GCP16 constitute a human protein fatty acyltransferase with specificity for H-and N-Ras. J. Biol. Chem. 280, 31141–31148 (2005).

21. Malgapo, M. I. P. & Linder, M. E. Substrate recruitment by zDHHC protein acyltransferases. Open Biol. 11, 210026 (2021).

22. Hong, J. Y. et al. High-Throughput Enzyme Assay for Screening Inhibitors of the ZDHHC3/7/20 Acyltransferases. ACS Chem. Biol. 16, 1318–1324 (2021).

23. Halbrook, C. J., Lyssiotis, C. A., Pasca di Magliano, M. & Maitra, A. Pancreatic cancer: Advances and challenges. Cell 186, 1729–1754 (2023).

24. Tang, Z. et al. GEPIA: a web server for cancer and normal gene expression profiling and interactive analyses. Nucleic Acids Res. 45, W98–W102 (2017).

25. Uhlén, M. et al. Tissue-based map of the human proteome. Science 347, 1260419 (2015).

26. Zhang, M. et al. A STAT3 palmitoylation cycle promotes TH17 differentiation and colitis. Nature 586, 434–439 (2020).

27. Liu, Y. et al. MAVS Cys508 palmitoylation promotes its aggregation on the mitochondrial outer membrane and antiviral innate immunity. Proc Natl Acad Sci USA 121, e2403392121 (2024).

28. Hancock, J. F., Magee, A. I., Childs, J. E. & Marshall, C. J. All ras proteins are polyisoprenylated but only some are palmitoylated. Cell 57, 1167–1177 (1989).

29. Roy, S. et al. Individual palmitoyl residues serve distinct roles in H-ras trafficking, microlocalization, and signaling. Mol. Cell. Biol. 25, 6722–6733 (2005).

30. Rocks, O. et al. An acylation cycle regulates localization and activity of palmitoylated Ras isoforms. Science 307, 1746–1752 (2005).

31. Cuiffo, B. & Ren, R. Palmitoylation of oncogenic NRAS is essential for leukemogenesis. Blood 115, 3598–3605 (2010).

32. Dougherty, M. K. et al. Regulation of Raf-1 by direct feedback phosphorylation. Mol. Cell 17, 215–224 (2005).

33. Rushworth, L. K., Hindley, A. D., O’Neill, E. & Kolch, W. Regulation and role of Raf-1/B-Raf heterodimerization. Mol. Cell. Biol. 26, 2262–2272 (2006).

34. Ritt, D. A., Monson, D. M., Specht, S. I. & Morrison, D. K. Impact of feedback phosphorylation and Raf heterodimerization on normal and mutant B-Raf signaling. Mol. Cell. Biol. 30, 806–819 (2010).

35. Amendola, C. R. et al. KRAS4A directly regulates hexokinase 1. Nature 576, 482–486 (2019).

36. Arora, N., Mu, H., Liang, H., Zhao, W. & Zhou, Y. RAS G-domains allosterically contribute to the recognition of lipid headgroups and acyl chains. J. Cell Biol. 223, (2024).

37. Weise, K., Triola, G., Brunsveld, L., Waldmann, H. & Winter, R. Influence of the lipidation motif on the partitioning and association of N-Ras in model membrane subdomains. J. Am. Chem. Soc. 131, 1557–1564 (2009).

38. Tran, N. H., Wu, X. & Frost, J. A. B-Raf and Raf-1 are regulated by distinct autoregulatory mechanisms. J. Biol. Chem. 280, 16244–16253 (2005).

39. Spencer-Smith, R. et al. RASopathy mutations provide functional insight into the BRAF cysteine-rich domain and reveal the importance of autoinhibition in BRAF regulation. Mol. Cell 82, 4262–4276.e5 (2022).

40. Nussinov, R., Tsai, C.-J. & Jang, H. Is Nanoclustering essential for all oncogenic KRas pathways? Can it explain why wild-type KRas can inhibit its oncogenic variant? Semin. Cancer Biol. 54, 114–120 (2019).

41. Liau, N. P. D. et al. Structural basis for SHOC2 modulation of RAS signalling. Nature 609, 400–407 (2022).

42. Kovrigina, E. A., Galiakhmetov, A. R. & Kovrigin, E. L. The ras G domain lacks the intrinsic propensity to form dimers. Biophys. J. 109, 1000–1008 (2015).

43. Lin, C.-C. et al. Receptor tyrosine kinases regulate signal transduction through a liquid-liquid phase separated state. Mol. Cell 82, 1089–1106.e12 (2022).

44. Zhou, M., Shi, N. & Li, P. Optogenetic control of biomolecular organization reveals distinct roles of phase separation in RTK signaling. Cell Chem. Biol. 32, 1503–1516.e5 (2025).

45. Lin, C.-W. et al. A two-component protein condensate of the EGFR cytoplasmic tail and Grb2 regulates Ras activation by SOS at the membrane. Proc Natl Acad Sci USA 119, e2122531119 (2022).

46. Ostrem, J. M., Peters, U., Sos, M. L., Wells, J. A. & Shokat, K. M. K-Ras(G12C) inhibitors allosterically control GTP affinity and effector interactions. Nature 503, 548–551 (2013).

47. Chen, W.-C. et al. Targeting KRAS4A splicing through the RBM39/DCAF15 pathway inhibits cancer stem cells. Nat. Commun. 12, 4288 (2021).

48. Kochen Rossi, J., et al. The differential interactomes of the KRAS splice variants identify BIRC6 as a ubiquitin ligase for KRAS4A. Cell Rep. 44, 115087 (2025).

49. Corcoran, R. B. et al. STAT3 plays a critical role in KRAS-induced pancreatic tumorigenesis. Cancer Res. 71, 5020–5029 (2011).

50. D’Amico, S. et al. STAT3 sustains tumorigenicity following mutant KRAS ablation. EMBO Rep. 26, 4900–4922 (2025).

51. Sadozai, H. et al. High hypoxia status in pancreatic cancer is associated with multiple hallmarks of an immunosuppressive tumor microenvironment. Front. Immunol. 15, 1360629 (2024).

52. Zhang, M. et al. SMAD2 S-palmitoylation promotes its linker region phosphorylation and TH17 cell differentiation in a mouse model of multiple sclerosis. Sci. Signal. 18, eadr2008 (2025).

53. Hurst, C. H., Turnbull, D. & Hemsley, P. A. Determination of Protein S-Acylation State by Enhanced Acyl-Switch Methods. Methods Mol. Biol. 2009, 3–11 (2019).

54. Zhou, Y. et al. SIGNAL TRANSDUCTION. Membrane potential modulates plasma membrane phospholipid dynamics and K-Ras signaling. Science 349, 873–876 (2015).

55. Zhou, Y. et al. Lipid-Sorting Specificity Encoded in K-Ras Membrane Anchor Regulates Signal Output. Cell 168, 239–251.e16 (2017).

56. Stirling, D. R. et al. CellProfiler 4: improvements in speed, utility and usability. BMC Bioinformatics 22, 433 (2021).

